# Open-science discovery of DNDI-6510, a compound that addresses genotoxic and metabolic liabilities of the COVID Moonshot SARS-CoV-2 Mpro lead inhibitor

**DOI:** 10.1101/2025.06.16.660018

**Authors:** Ed J. Griffen, Daren Fearon, Briana L. McGovern, Lizbe Koekemoer, Blake H. Balcomb, Tamas Szommer, Gwendolyn Fate, Ralph P. Robinson, Bruce A. Lefker, Shirly Duberstein, Noa Lahav, The COVID Moonshot Consortium, Stephanie Braillard, Laura Vangeel, Manon Laporte, Fabienne Burgat Charvillon, A. Kenneth MacLeod, Andrew Wells, Pauline Garner, Richard Knight, Paul Rees, Anthony Simon, Dirk Jochmans, Johan Neyts, Kevin D. Read, Haim Barr, Matthew Robinson, Alpha Lee, Nir London, John Chodera, Frank von Delft, Kris M. White, Ben Perry, Peter Sjö, Annette von Delft

## Abstract

The 2020 SARS-CoV-2 coronavirus pandemic highlighted the urgent need for novel small molecule antiviral drugs. (S)-**x38** DNDI-6510 is a non-covalent SARS-CoV-2 main protease inhibitor developed by the open science collaboration COVID Moonshot.

Here, we report on the metabolic and toxicologic optimization of the lead series previously disclosed by the COVID Moonshot Initiative, leading up to the selection of (S)-**x38** DNDI-6510 as the preclinical candidate. We describe the thorough profiling of the series, identifying key risks such as formation of genotoxic metabolites and high clearance, which were successfully addressed during lead optimization. In addition, we disclose the *in vitro* and *in vivo* evaluation of (S)-**x38** DNDI-6510 in pharmacokinetic and pharmacodynamic models, exploring multiple approaches to ameliorate rodent-specific metabolic clearance, and show that both co-dosing of (S)-**x38** DNDI-6510 with an ABT inhibitor and utilizing a metabolically humanized mouse model (8HUM) achieve significant improvements in exposure. Through comparisons of ABT co-dosing and humanized mouse models in efficacy experiments, we demonstrate that continuous exposure over cellular EC_90_ is required for SARS-CoV-2 antiviral efficacy *in vivo* in an antiviral model using a mouse-adapted SARS-CoV-2 strain. Finally, (S)-**x38** DNDI-6510 was assessed in maximum tolerated dose experiments in two species, demonstrating significant in vivo PXR-linked auto-induction of metabolism, leading to the discontinuation of this compound.

In summary, we report the successful effort to overcome series-specific AMES liabilities in a lead development program. Downstream optimization of existing series will require in-depth optimization of rodent-specific liabilities and metabolic induction profile.

## Introduction

The emergence of COVID-19 as a global pandemic clearly highlighted the fundamental lack of access to safe, efficacious, globally accessible, and affordable broad-spectrum oral antivirals. SARS-CoV-2 has caused over 7 million confirmed deaths to date, with the estimated number exceeding 28 million^1^.

Despite declining disease severity and number of reported cases, as well as access to effective vaccinations, coronaviruses will continue to present a significant threat to global health for the foreseeable future. In addition, chronic manifestations of viral infection i.e. long COVID, estimated to affect up to 10% of infected individuals, remain untreated^2^.

Through impressive efforts novel SARS-CoV-2 antivirals such as nirmatrelvir and ensitrelvir were developed in record time^3,4^, however, global access to these drugs is still limited^5^. Follow-on compounds from the groups at Pfizer and Shionogi address key liabilities of the first generation inhibitors, such as ibuzatrelvir^6^, which avoids the need for ritonavir that was required for Paxlovid (nirmatrelvir plus ritonavir)^7^, and S-892216^8^, which is now being assessed in Phase 2 clinical trials as a long-acting alternative to ensitrelvir intended for prophylactic treatment. Of note, ensitrelvir was also shown to prevent transmission of SARS-CoV-2 infection in a post exposure prophylaxis trial^9^.

Case reports on increasing levels of SARS-CoV-2 drug resistance for mutants^10,11^ further underline the need for additional antiviral treatment options. Since COVID-19 was not an isolated event, but rather the latest example in a series of viral pandemic threats to human health^12^, continued antiviral drug discovery and development needs to be a key pillar of ongoing pandemic preparedness efforts.

The COVID Moonshot was an unprecedented open-science effort, based on a crystallographic fragment screen, that led to the identification and initial optimization of an isoquinoline-based series of SARS-CoV-2 main protease (MPro) inhibitors^13^. The Moonshot lead (S)-x1, named MAT-POS-e194df51-1 in the original publication^13^, is a potent and selective inhibitor of SARS-CoV-2 Mpro activity, which translates into potent antiviral activity against SARS-CoV-2 variants of concern *in vitro* (Figure 1).

**Figure 1.**
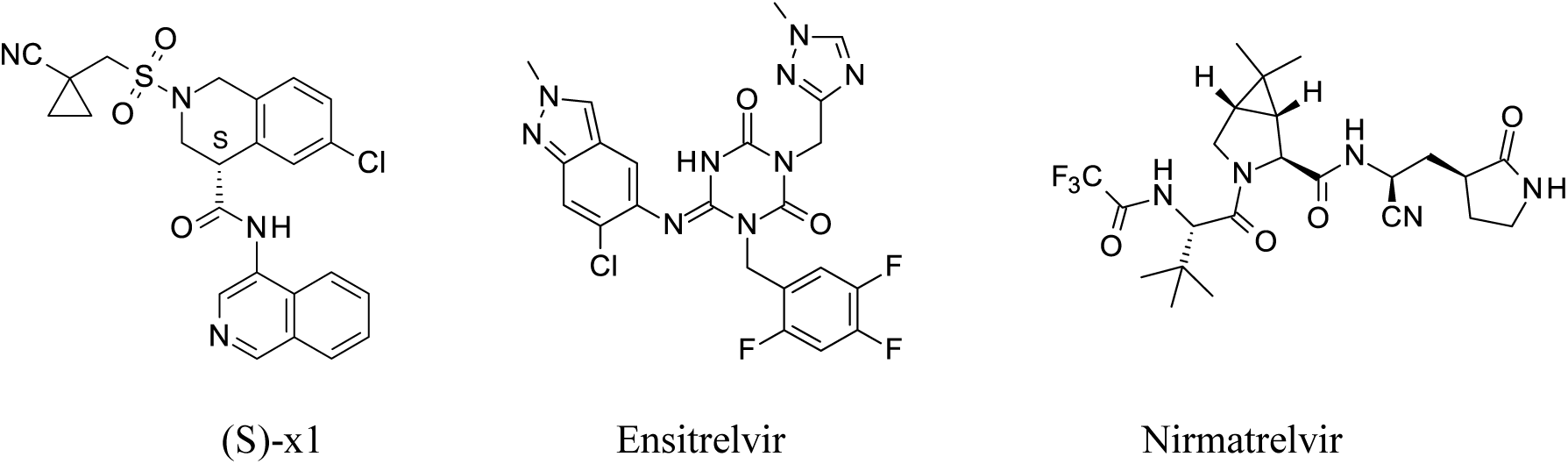
Chemical structures of the SARS-CoV-2 Mpro inhibitors (S)-x1, ensitrelvir, and nirmatrelvir

The structural basis for the interaction of (S)-x1 with the SARS-CoV-2 Mpro is well characterized, with ligand-protein crystal structures confirming the importance of the isoquinoline to His163 H-bond in the P1 pocket, and the deep insertion of the aromatic -Cl in the P2 binding pocket^13^. Further, structural data also confirmed the non-covalent binding mode for this series^13^. The COVID Moonshot lead compound (S)-x1 showed a well-balanced property profile, with good solubility and clean protease selectivity, and no major flags were detected in broad selectivity panel testing^13^. However, a number of potential liabilities were identified for (S)-x1. An early literature screen for structural alerts highlighted a potential risk for formation of genotoxic metabolites associated with the embedded amino isoquinoline fragment^14^, later confirmed in AMES assays. In addition, *in vitro* pharmacokinetic data indicated the need for an improvement of the metabolic clearance profile of the series and to abrogate the potential for racemization of the one stereocenter.

Here, we describe the step-wise process taken to remove of the genotoxic and metabolic liabilities of (S)-x1, leading to the discovery of the optimized lead candidate (S)-**x38,** also named DNDI-6510, a non-covalent, potent SARS-CoV-2, isoquinoline-based MPro inhibitor. (S)-**x38** was extensively profiled in *in vitro* and *in vivo* assays, and demonstrated potent antiviral activity across a range of SARS-CoV-2 variants of concern. Based on this data, (S)-**x38** was advanced to preclinical toxicology studies, where rodent-specific PXR-linked induction was identified preventing full toxicological assessment of the compound at relevant plasma levels leading to the decision to discontinue the preclinical development of (S)-**x38**. In line with the COVID Moonshot open-science principles, all discovery data, including antiviral, pharmacokinetic and safety profiling generated during the lead-optimization phase useful for the research community are being disclosed through an open CDD vault and a ChEMBL deposition.

## Results

### Optimization of Ames profile/ isoquinoline mutagenicity

As a first step towards addressing the key liabilities of the isoquinoline-based inhibitors, we focused on addressing the genotoxicity risk indicated by the Ames activity observed for a number of (S)-x1 structural analogues. Varying the substitution in the 4-amino isoquinoline 7-position, offers a low probability of resolving the Ames liability (Figure 2b). These results correlate well with prediction of their metabolic activation^15^.

**Figure 2.**
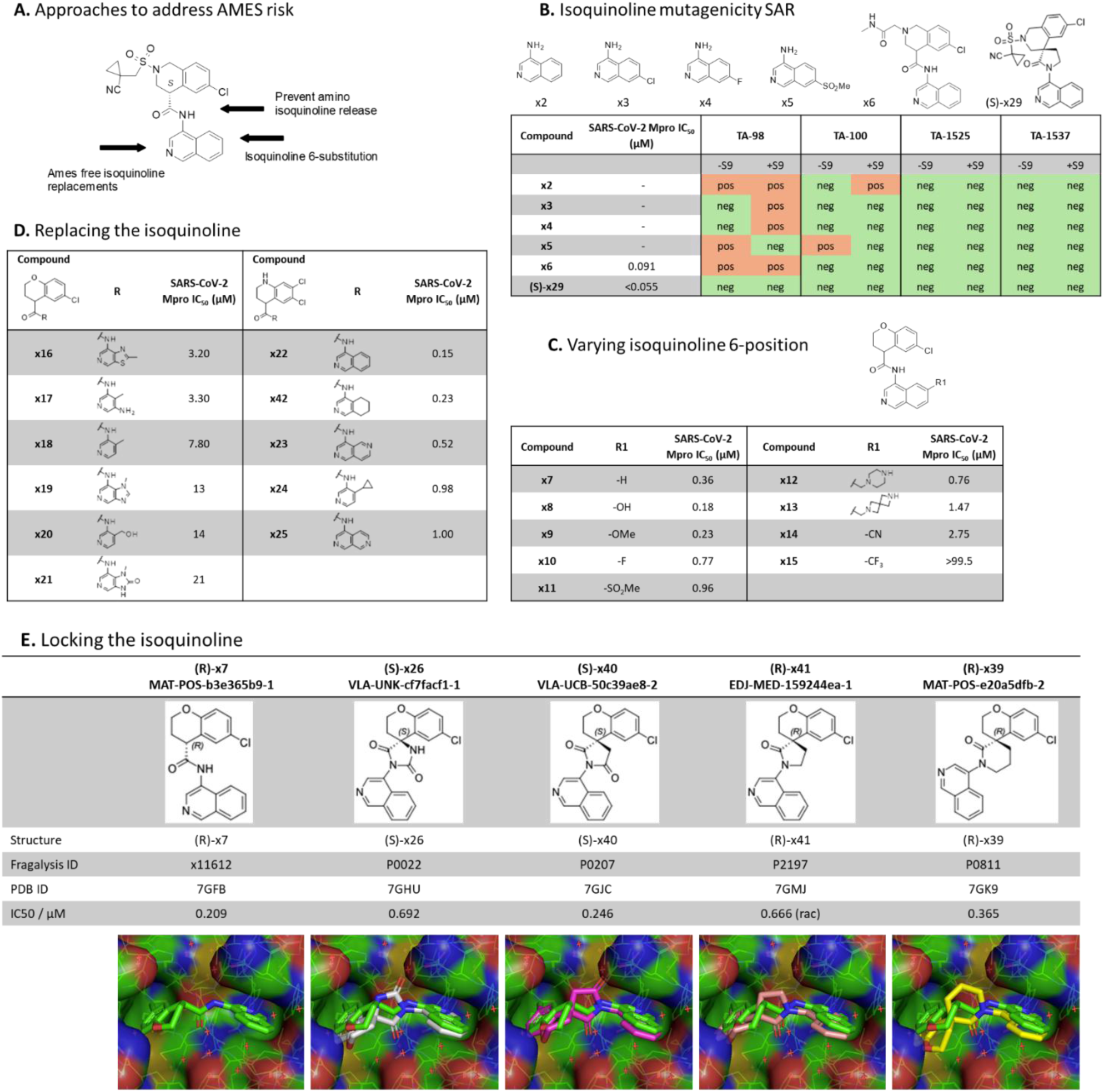
Solving the AMES liability (A) Conceptual approaches to address the Ames risk are shown. (B) Mutagenicity of a range of isoquinolines in a range of Salmonella typhimurium (TA) strains used to detect mutagenic compounds, with or without metabolic activation derived from rat liver microsomes (S9) to mimic *in vivo* metabolism. (C) Approach of 6-substituted isoquinolines to avoid Ames activity. (D) Impact of different isoquinoline replacements on SARS-CoV-2 MPro activity. (E) Cyclisation strategies to “lock-in” the amino isoquinoline avoiding formation of Ames active metabolites.

Instead, we focused our efforts on identifying an Ames-free core using three approaches (Figure 2a): 1) substitution to obtain Ames free 4-amino isoquinolines by investigating different substituents in the isoquinoline 6-position (Figure 2c); 2) replacing the isoquinoline with amino heterocycles with a reduced mutagenicity risk maintaining the critical isoquinoline His163 H-bond interaction (Figure 2d); and 3) spirocyclisation preventing release of potential Ames active metabolites (Figure 2e).

In the first approach (Figure 2c), assuming that the Ames activity was due to nitrosamines conjugating with DNA caused by oxidation of the amine via the hydroxylamine^16^, we hypothesized that electron withdrawing substituents at the 6 position would reduce the rate of oxidation. Unfortunately, the electron withdrawing substituents (F, SO_2_Me, CN, CF_3_) were less active as Mpro inhibitors, while the electron donating substituents (OH, OMe) were more potent protease inhibitors, we assume due to better H-bonding to the His163. We hypothesized that (i) the amino methyl substituents are protonated at physiological pH and therefore act as electron withdrawing substituents, again reducing Mpro activity; and (ii) that more electron rich amino heterocycles would oxidize more readily to nitrosamines; therefore, further exploration of this approach was discontinued.

In the second approach, we synthesized a series of isoquinoline replacements (Figure 2d). Most replacements demonstrated a significant (>3-fold) drop in SARS-CoV-2 Mpro potency, with only the tetrahydroisoquinoline **x42** able to maintain potency similar to the original isoquinoline **x22**. However, we considered the significant risk for oxidative re-aromatization back to the isoquinoline to be too high and decided to discontinue this exploration.

In the third approach, also explored in the discovery of potent hydantoin-based other SAR-CoV-2 Mpro inhibitors^17^, we synthesized [5,6]-spirocyclic matched pair analogues of the amide-linked isoquinoline-based inhibitors, expected to have a good overlap with available X-ray structures. Although the X-ray structures of the first hydantoin based derivatives align well their potency did not appear particularly promising (Figure 2e). Further exploration of imide- and lactam-based spirocycles showed a lower drop in potency, which we hypothesized could be attributed to progressively less desolvation upon binding. The matched-pair examples prepared clearly demonstrated the viability of this approach in retaining potent SARS-CoV-2 Mpro activity, Table 1. In particular, even though there was a 3-fold loss of MPro potency observed for matched pairs x7/x26 and x27/x28/x29, the glycine amide substituent x6/x31 was equipotent (Table 1). We also explored a six membered ring lactam, hoping that the additional lipophilicity would provide more potency or that the slight difference in conformation could be beneficial, however, it was significantly less potent, with the X-ray crystal structure demonstrating that this sub-structure caused a significant movement of the ligand in the active site comparing with the non-cyclized ligand **x7** (Figure 2e). The active enantiomers of (*S*)-**x29,** as well as later optimized leads (*S*)-**x31,** (*S*)-**x37** and (*S*)-**x38** did not show AMES activity when tested against the TA-98 and T-100 strains with or without the presence of liver S9 fraction metabolizing enzymes (Supplementary Table 1). In addition, the creation of an all-carbon substituted stereocenter removed any risk for racemization. Since the lactam-based series showed a low drop in Mpro potency upon moving to the conformationally locked system, it was selected for further elaboration.

**Table 1.**
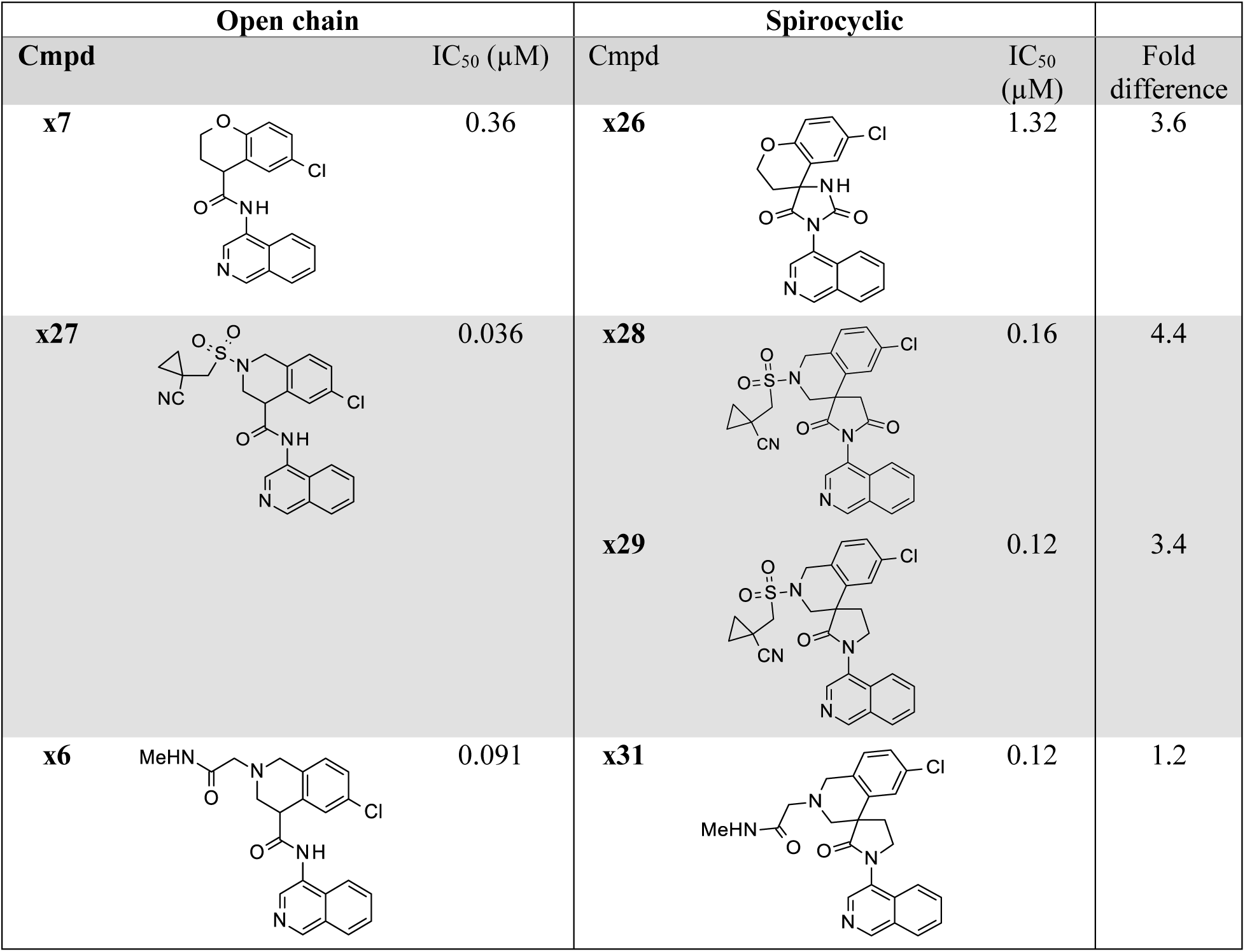
SARS-CoV-2 Mpro activity for N-captured isoquinoline open chain and spirocyclic matched pairs.

| Open chain |  | Spirocyclic |  | Fold difference |
| --- | --- | --- | --- | --- |
| Cmpd | IC<sub>50</sub> (μM) | Cmpd | IC<sub>50</sub> (μM) |  |
| **x7** | 0.36 | **x26** | 1.32 | 3.6 |
| **x27** | 0.036 | **x28** | 0.16 | 4.4 |
|  |  | **x29** | 0.12 | 3.4 |
| **x6** | 0.091 | **x31** | 0.12 | 1.2 |

### Reduction of metabolism

In parallel to addressing the AMES genotoxicity risk, we addressed the metabolic profile of the series, whilst retaining highly potent SARS-CoV-2 Mpro inhibitors. Through a systematic evaluation of compound **x6** we identified metabolically exposed positions including possible amide bond cleavages, isoquinoline N-oxidation and aromatic oxidation, and benzylic oxidation of the tetrahydroisoquinoline moiety (Figure 3a). Recognizing that initial examples of the series showed rapid in-vivo clearance in rodent species, without always a clear connection to in-vitro metabolic stability, we chose to monitor the composite effect on clearance using i.v. cassette dosing in rats.

**Figure 3.**
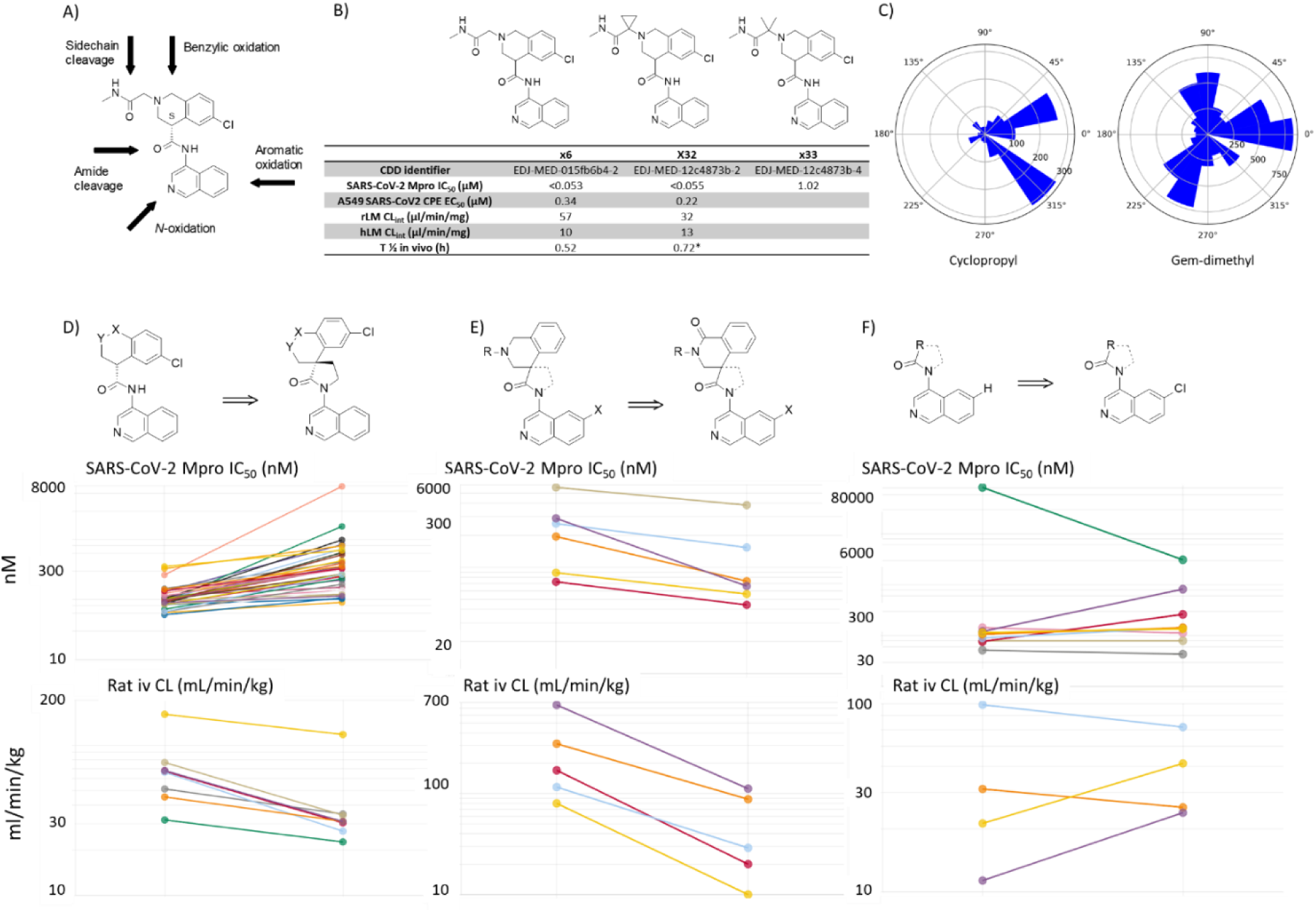
Overview of approaches taken to address metabolism. (A) Conceptional overview of potential metabolic liabilities of the COVID Moonshot lead (S)-x1, including sidechain cleavage, benzylic oxidation, amide cleavage, as well as isoquinoline *N*-oxidation and aromatic oxidation. (B) Approaches to address potential sidechain cleavage with effects on SARS-CoV-2 enzyme potency and rat and human liver microsome clearance in vitro are depicted. (C) Polar histogram plots in Conquest (Cambridge Crystallographic Data Centre (CCDC)^18^ of carbonyl-methylene angle distribution plots for acyl cyclopropyl compounds (n=1347) (left) and carbonyl-methyl angle distribution for acyl gem-dimethyl compounds (n=6433) (right). (D) Aminoquinoline cyclisation to address potential amide cleavage. The effect in a matched pair analysis (not cyclized, left) and (cyclized, right) on Mpro IC50 (top) and Rat IV clearance (bottom panel) is depicted. (E) Effect of benzylic oxidation in a matched pair analysis (tetrahydroisoquinoline, left) and (tetrahydroisoquinolone, right) on Mpro IC50 (top) and Rat IV clearance (bottom panel) is depicted. (F) Effect of blocking the 6-isoquinoline position in a matched pair analysis (6-H, left) and (6-Cl, right) on Mpro IC50 (top) and Rat IV clearance (bottom panel) is depicted.

The risk for amide sidechain cleavage was addressed by increasing steric hindrance to reduce the rate of attack on the amide carbonyl by gem-dimethylation and cyclopropanation of the alpha-carbon. Whilst enzyme potency is maintained with cyclopropanation, it is lost with gem-dimethylation (Figure 3b). Further, beyond maintaining potency, the cyclopropyl derivative indeed shows improved clearance in vitro in rat liver microsomes, as well as an improved in vivo half-life (Figure 3b). This may be explained by a difference in conformational preference for the gem-dimethyl vs. cyclopropyl substituted sidechains. An analysis of alpha-substituted gem-dimethyl and cyclopropyl amide analogues found in the collection of the Cambridge Crystallographic Data Centre clearly indicates a conformational preference for the cyclopropyl to be orientated *cis* in relation to the carbonyl group, whilst the gem-dimethyl substituted sidechain is preferentially orientated *trans* (Figure 3c)^18^.

We rationalized that, with the formation of a spirocyclic ring, the central amide bond would be stabilized and less likely to be hydrolyzed (Figure 3d). This was indeed found to be the case when comparing 10 matched pair analogues, an average decrease in rat *in vivo* CL_int_ of a factor of 1.6 was observed (Figure 3d, bottom panel), whilst a median loss in Mpro potency of a factor of 2.5 was detected when comparing 28 matched pair examples (Figure 3d, top panel). Blocking the tetrahydroisoquinoline benzylic position by the introduction of a carbonyl group led to a significant improvement of the rat *in vivo* clearance by an average factor of 6.3 in 6 matched pairs (Figure 3e, bottom panel), which was accompanied by an improvement of Mpro potency by a factor of 1.7 (n = 6) (Figure 3e, top panel). Finally, blocking isoquinoline aromatic oxidation by introducing Cl (or F) in the 6- or 7-position had an overall neutral effect on potency (n=7) and stability (n=4) (Figure 3f), with F substitution in the 6-position having an overall negative effect on *in vivo* clearance (Figure 3f); this approach was thus not pursued further. In the COVID Moonshot and throughout this lead optimization project, continuous structural assessment was a cornerstone of the medicinal chemistry strategy^13,19^. The campaign leveraged high-throughput X-ray crystallography to generate nearly 500 ligand-bound protein structures, enabling real-time feedback on compound binding modes^20^. This structural data was integrated with biochemical assays and computational predictions to inform compound design and prioritization. During this period a large number of other medicinal chemistry opportunities were considered. As we were generating protein-ligand crystal structures in parallel to determining enzyme inhibition potencies, we were rapidly able to prune our approaches, avoiding preparation of molecules where the probability of productive binding was low.

### Selection and in vitro profiling of lead candidates

Our optimization efforts identified three potent SARS-CoV-2 Mpro inhibitors for further evaluation (Figure 4a). All three compounds had potent antiviral activity in A549 SARS-CoV-2 infected cells (Figure 4c), promising physiochemical properties (Table S3), and were negative in Ames and in vitro micronucleus genotoxicity assays (Table S1). Plasma protein binding measurements determined high unbound fractions across species for all three compounds (Table S4). In particular, the fraction unbound for (S)-x38 was 54%, 12% and 46% in rat, dog and human plasma, respectively (Table S4). Further, all lead compounds demonstrated a favorable selectivity profile in a protease screen evaluating several proteases including caspases 1 to 9, cathepsin B, K, L, S and calpain 1 (Table S5), and no flags in a broad panel of G-protein coupled receptors, transporters, kinases and phosphodiesterase screens (Table S6). In addition, no activity against cardiac ion channels hERG, Kv1.1, Cav1.2, and Nav1.5 was detected up to concentrations of 30 uM (Table S7).

**Figure 4:**
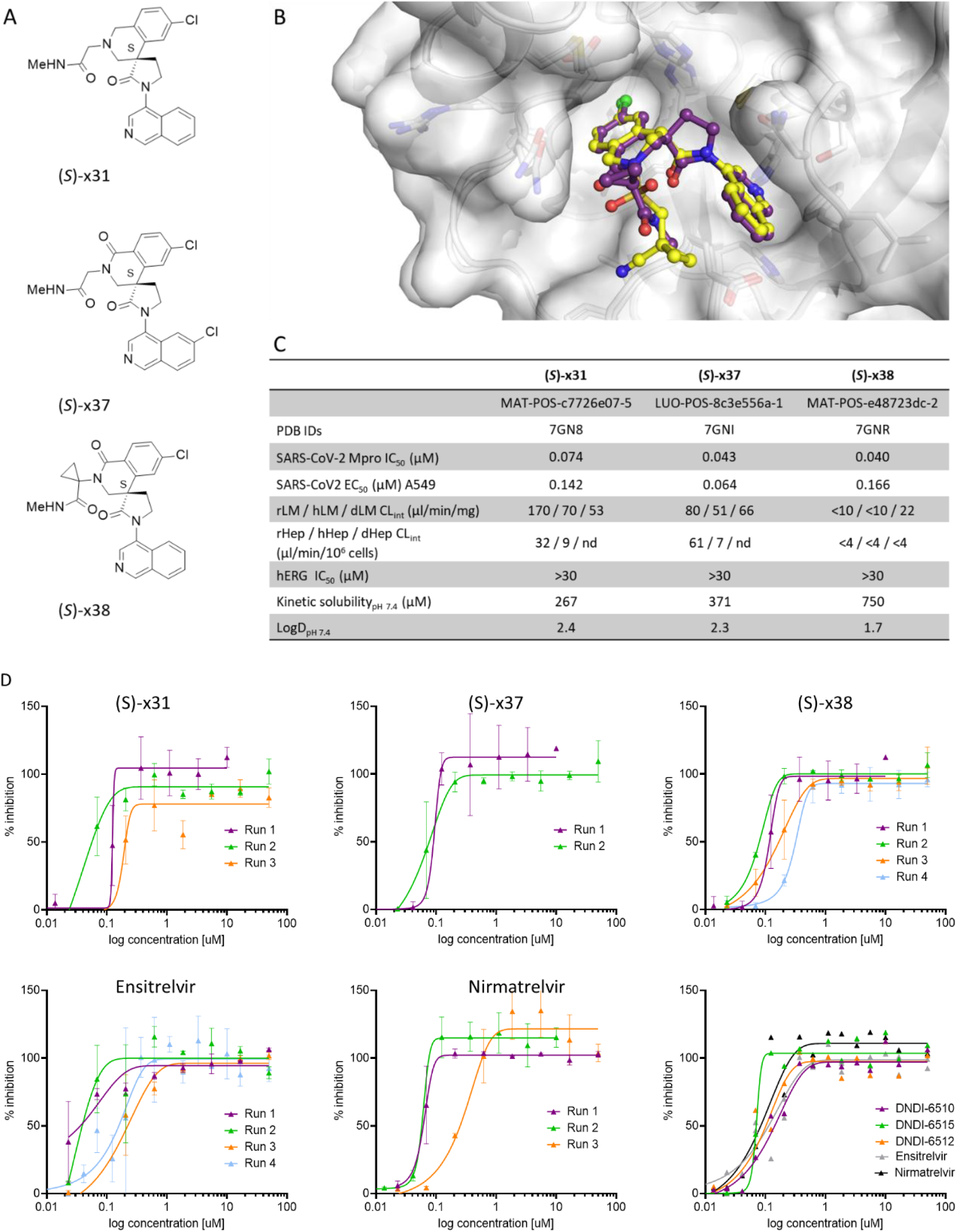
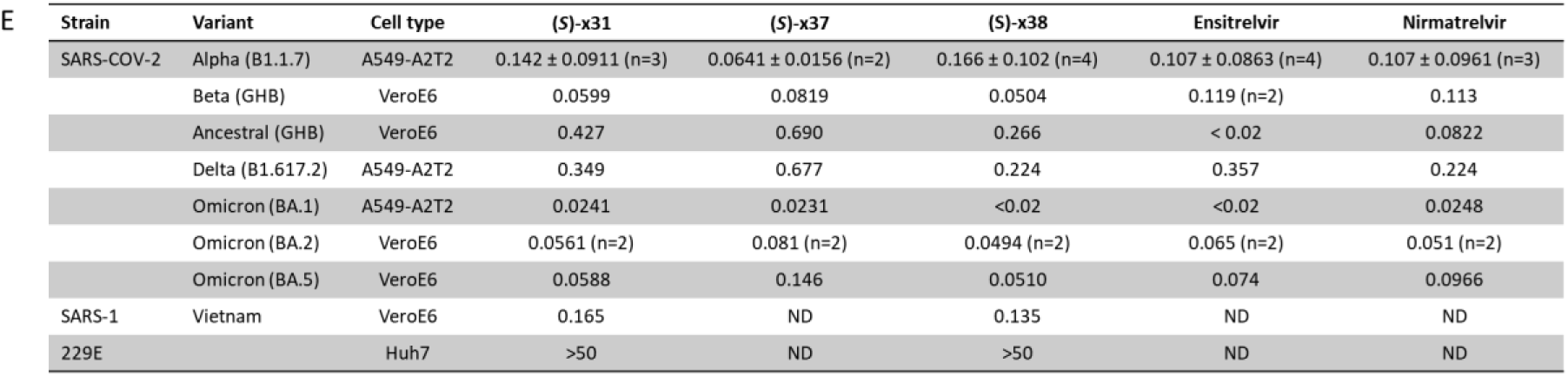
Identification and characterization of lead candidates with ameliorated AMES risk. (A) Structure of lead candidates. (B) In vitro characterization of the three lead candidates, with potency, solubility, metabolic and safety parameters detailed. (C) Overlay of co-crystal structures of COVID Moonshot lead compound (S)-x1 (yellow, PDB ID: 7GN8) and (S)-x38 (purple, PDB ID: 7GNR), demonstrating the preservation of key molecular interactions in S1 and S2 pockets are maintained. (C) Comparison of lead compounds (S)-31, (S)-37 and (S)-38 as well as ensitrelvir and nirmatrelvir in an *in vitro* SARS-CoV-2 cellular antiviral efficacy in A549 cells overexpressing ACE-2 and TMPRSS-2. (E) Cross-reactivity of lead compounds (S)-31, (S)-37 and (S)-38 and controls ensitrelvir and nirmatrelvir in cellular antiviral assays of SARS-CoV-2 variants of concern, as well as SARS-CoV-1 and 229E.

Continuous, real-time structural analysis affirmed that recapitulating key molecular interactions identified in the original crystallographic fragment screen^20^ was essential for driving potency during early hit-to-lead development and lead optimization, particularly with the spirocyclic lactam series. Specifically, a conserved hydrogen bond between the isoquinoline nitrogen and His163 in the S1 pocket, a hydrogen bond between a carbonyl group and the backbone NH of Glu166, and hydrophobic interactions involving an aromatic chlorine atom in the S2 pocket were consistently associated with high-affinity binding. The crystal structure of (S)-x38 (Figure 4b, PDB ID: 7GNR) confirms the preservation of all three interactions, underscoring their importance in the design of potent inhibitors. The cross-species metabolic stability in *in vitro* microsomal (Table S8) and hepatocyte (Table S9) assays highlighted the favorable stability profile for compound (S)-**x38** (Figure 4b), which was thus further profiled in *in vivo* pharmacokinetic (PK) and efficacy studies.

The three optimized lead compounds showed potent antiviral efficacy against SARS-CoV-2 in a cellular A549 assay comparable to both ensitrelvir and nirmatrelvir, with antiviral cellular EC_90_ of 142 nm, 64nM and 66 nM for (S)-**x31**, (S)-**x37** and (S)-**x38**, respectively (Figure 4d). No cytotoxicity was detected in a number of cell lines evaluated (Table S10). Furthermore, the compound was active against known SARS-CoV-2 variants of concern (VOC) with potencies ranging from EC_50_ of less than 20 nM (Omicron BA.1) to 224 nM against the delta strain (Figure 4e). When tested against other coronaviruses in a MPro enzyme panel, (S)-x38 was active against SARS-COV-1, showed weak activity against other coronaviruses OC34 (IC_50_ 7.78 µM) and HKU1 (IC_50_ 5.73 µM), but showed no activity against MERS, 229E, or NL63 (IC_50_ >20 µM) in biochemical assays (Table S11) and against 229E in cellular assays (Figure 4e).

### In vivo profiling of (S)-x38

With the selected lead (S)-**x38**, a significant improvement in metabolic stability was achieved, with RLM apparent intrinsic clearance decreasing from 605 µl/min/mg for (S)-x1 (Moonshot lead) to 16 µl/min/mg ((S)-**x38**) (Figure 5a). However, despite the significant improvements in metabolic clearance, the series retained significant rodent-specific metabolic liabilities, with a high clearance of 90 mL/min/kg in wild-type mice upon oral dosing (Table S12). Two approaches were investigated to enable efficacy experiments in a mouse model; (1) co-dosing with 1-aminobenzotriazole (ABT), a broad spectrum CYP inhibitor, and (2) a humanized mouse model (8HUM) as previously described^21^.

**Figure 5:**
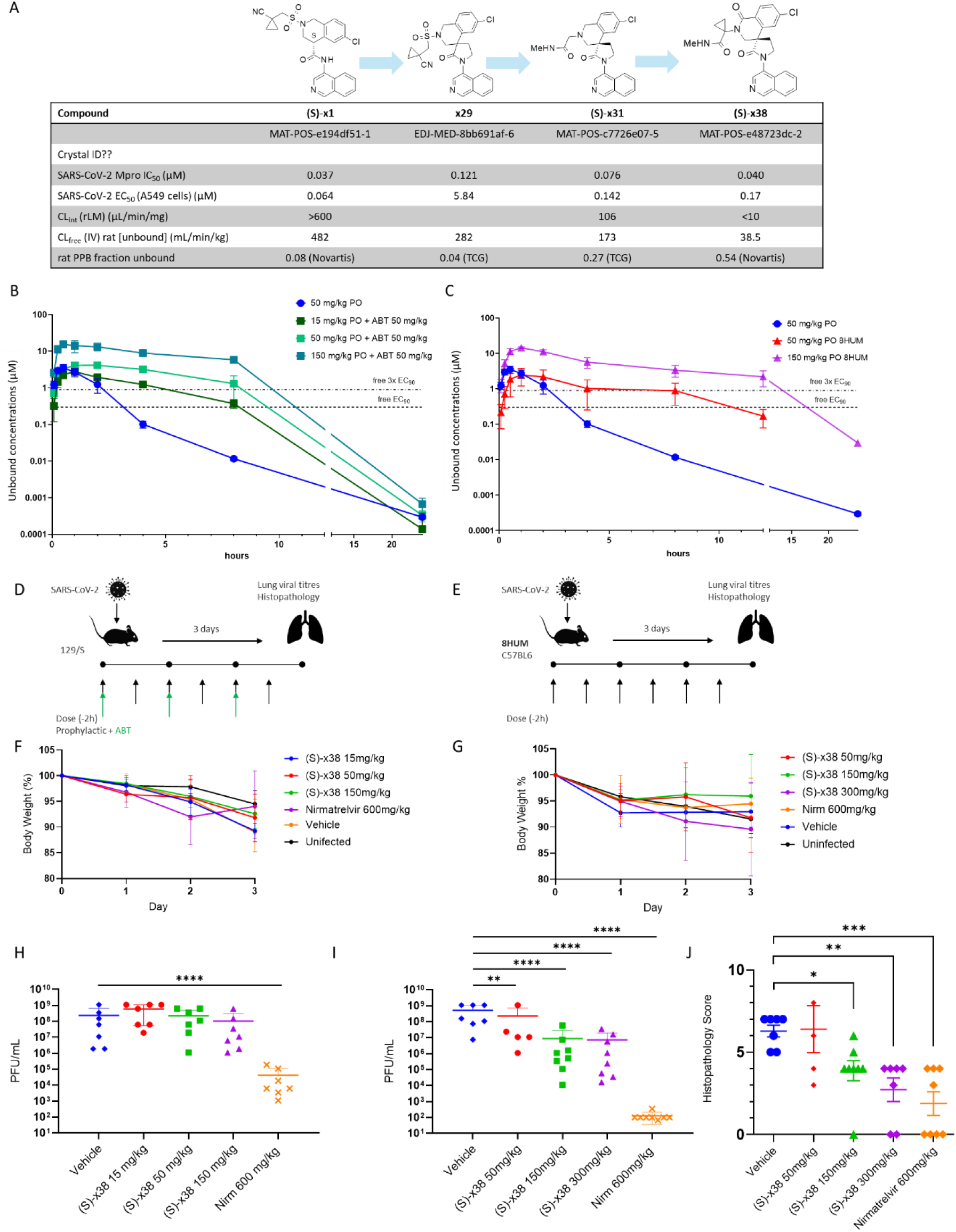
(S)-x38 in vivo pharmacokinetic profiling and animal efficacy in SARS-CoV-2 infected mice. (A) Overview on characteristics of key compounds explored during lead optimization, highlighting the progress made on optimizing rodent metabolism. (B) Pharmacokinetic profiling of orally dosed ASAP-0017445 in wild-type mice (50 mg/kg), ASAP-0017445 in wild-type mice co-dosed with 1-aminobenzotriazole (ABT) (15 mg/kg, 50 mg/kg, 150 mg/kg), and (C) ASAP-0017445 dosed in humanized mice (8HUM) (50 mg/kg, 150 mg/kg). (D) Experimental layout of the ABT co-dosing SARS-CoV-2 efficacy experiment. (E) Experimental layout of the SARS-CoV-2 efficacy experiment in 8HUM mice. For both groups, mice were challenged intranasally with 2.5 × 10^4^ PFU of mouse-adapted SARS-CoV-2 (MaA10). Animals were orally administered various doses of (S)-x38 BID, with or without 50 mg/kg 1-aminobenzotriazole (ABT) co-dosing, or 600 mg/kg Nirmatrelvir BID, or vehicle (placebo) 2 hours before infection. Body weights were assessed daily for all groups, and are shown for (F) the ABT co-dosing experiment, and (G) the experiment in humanized (8HUM) mice. Animals were euthanized at 3 days post infection (dpi) and lungs collected for virus titers. Lung viral titers at day 3 post infection are shown for (H) the ABT co-dosing experiment, and (I) the experiment in humanized (8HUM) mice. (J) Effect of (S)-x38 on histopathology scores in a humanized mouse model is depicted. Data was log-transformed and analyzed by two-way ANOVA (*P < 0.05, **P < 0.01, ***P < 0.001, and ****P < 0.0001).

Co-dosing with ABT can significantly alter the metabolism of small molecules in rodents, with plasma clearance of metabolically cleared test compounds reduced by over 90% in rats at ABT doses of 50 mg/kg^22^. We measured *in vivo* PK in mice, co-dosing 15, 50, and 150 mg/kg of (S)-**x38** with a 50 mg/kg dose of ABT. ABT co-dosing reduced the observed oral clearance in wildtype animals from 90 mL/min/kg to 13 to 20 mL/min/kg, increasing free exposure durations over SARS-CoV-2 antiviral 3x EC90 from 2.3 h to 4.9 h (15 mg/kg dose), 8.6 h (50 mg/kg), and 11.3 h (150 mg/kg), respectively (Figure 5c, d, Table S12).

Following confirmation of relatively high metabolic stability in vitro with 8HUM hepatic microsomes (Table S8), we then used a humanized mouse model (8HUM) in which 33 mouse CYP enzymes, the mouse transcription factors Pregnane X Receptor (PXR), and the Constitutive Androstane Receptor (CAR) are replaced with human CYP enzymes, PXR and CAR to investigate the pharmacokinetic profile of (S)-**x38**^21^. (S)-**x38** showed reduced oral clearance in 8HUM (48 mL/min/kg at 50 mg/kg, 21 mL/min/kg at 150 mg/kg) in comparison to wildtype (90 ml/min/kg at 50 mg/kg), with plasma unbound exposure over 3x free SARS-CoV-2 EC90 of 8 h at 50 mg/kg and 14.4 hours at 150 mg/kg (Figure 5c, d, Table S12).

Next, both the ABT co-dosing model and the humanized mouse model 8HUM were used to perform animal efficacy experiments against SARS-CoV-2. (Figure 5d, e). No effects on body weight (Figure 5f), or reduction in lung viral loads was achieved for all three DNDI-6510 doses (15, 50 and 150 mg/kg PO), whilst a 4-log reduction in viral load was observed for the positive control nirmatrelvir at 600 mg/kg (Figure 5h). In contrast, a significant reduction in lung viral load measured by TCID50 was observed for all dosing groups with a dose dependent effect (Figure 5i), with similar reductions in histopathology scores observed for both (S)-x38 and positive control nirmatrelvir at 600 mg/kg (Figure 5j). These results are in line with the higher exposures observed in the humanised mouse model 8HUM (AUC_last_ (ng.h/mL) 7,125,320) compared to ABT co-dosing (AUC_last_ (ng.h/mL) 146,021) in the preceding PK studies. Furthermore, trough drug concentrations for (S)-**x38** and nirmatrelvir were obtained from satellite animals for each dosing group prior to the morning dose (D1 24h, D2 36h, D3 72h), which demonstrated higher exposures in 8HUM animals for both (S)-**x38** and nirmatrelvir, in line with viral load reductions observed in the lung (Figure S1).

### Maximum tolerated dose studies and PXR-linked induction

To select the second toxicology species, drug metabolizing enzymes and drug metabolites were studied. (S)-**x38** showed a low clearance in liver microsomes (Table S8) across preclinical species, confirming the successful optimization of metabolic liabilities of the series. In a cross-species comparison of *in vitro* metabolites in cryopreserved hepatocytes, human, rat and minipig show no measurable turnover (CL_int_ <4 µL/min/10^6 cells), whilst dog, monkey and mouse show low CL_int_-values up to 15 µL/10^6^ cells /min (Table S9). In spite of the low turnover, multiple oxidative metabolites and GSH adducts were detected, with LC-MS/MS analysis identifying nine major metabolites across species (Figure S2a). No major human metabolites were detected after 80 min incubation (parent 100%), with only traces of M7 and M9 detected (M7 −2; M9 +14). Both identified human metabolites were also detected in rat, mouse, and dog hepatocytes, solidifying the choice of dog as the second toxicological species (Figure S2b).

(S)-**x38** revealed no reversible inhibition (IC50 > 25 μM) of the major human CYP enzymes including CYP1A2, CYP2B6, CYP2C8, CYP2C9, CYP2C19, CYP2D6 and CYP3A4 in human liver microsomes (HLM) (Table S14). Further, (S)-**x38** did not cause time-dependent inhibition of the catalytic activity of the major human CYPs in HLM, with no IC50 shift observed after a 30 min preincubation of (S)-**x38** in HLM in the absence or presence of CYP cofactor NADPH (Table S14), and no induction observed for human PXR in initial screening measurements.

Assessment of pharmacokinetic properties of (*S*)-**x38** in two species, rat and dog, demonstrated good oral bioavailability of up to 37% in rat, and >100% in dog, with intravenous clearance of 25 ml/min/kg in rat and 23.5 ml/min/kg in dog (Table S13). Pharmacokinetic predictions using allometric scaling methods using data from two species resulted in human predictions of 1010 mg BID or 297 mg TID, targeting continuous exposure over free EC90 380 nM (Table S2), making (S)-**x38**, henceforth known as DNDI-6510, a viable preclinical candidate for translation.

Next, (*S*)-**x38** DNDI-6510 was evaluated in repeat oral dosing studies as part of a maximum tolerated dose study in two species. In rat, (S)-**x38** DNDI-6510 showed significantly reduced exposures on day 3 in comparison to day 1, in particular for the higher dosing groups 300 mg/kg (AUC_last_ D1 mean 51,567 ± 21,975 ng·h/ml vs D3 mean 13,865 ± 8330 ng·h/ml) and 1000 mg/kg (AUC_last_ D1 mean 193,833 ± 67182 ng·h/ml vs D3 mean 46,217 ± 33034 ng·h/ml) (Figure 6a). A follow-on mouse PK study over 5 days at 100, 300 and 1000 mg/kg in male animals only confirmed this observation, with exposures reduced across groups (100 mg/kg AUC_last_ D1 mean 25507.3 ± SD 3500.9 ng·h/ml vs D3 mean 3792.3 ± SD 324.4 ng·h/ml) (Figure 6b). In contrast, in a multi-day dog study at 100 mg/kg once daily, increased exposures on day 7 compared to day 1 were observed in both male and female animals (AUC_last_ D1 mean 62167 ± 21085 ng·h/ml vs D7 mean 222500 ± 90734 ng·h/ml) (Figure 6c, d).

**Figure 6:**
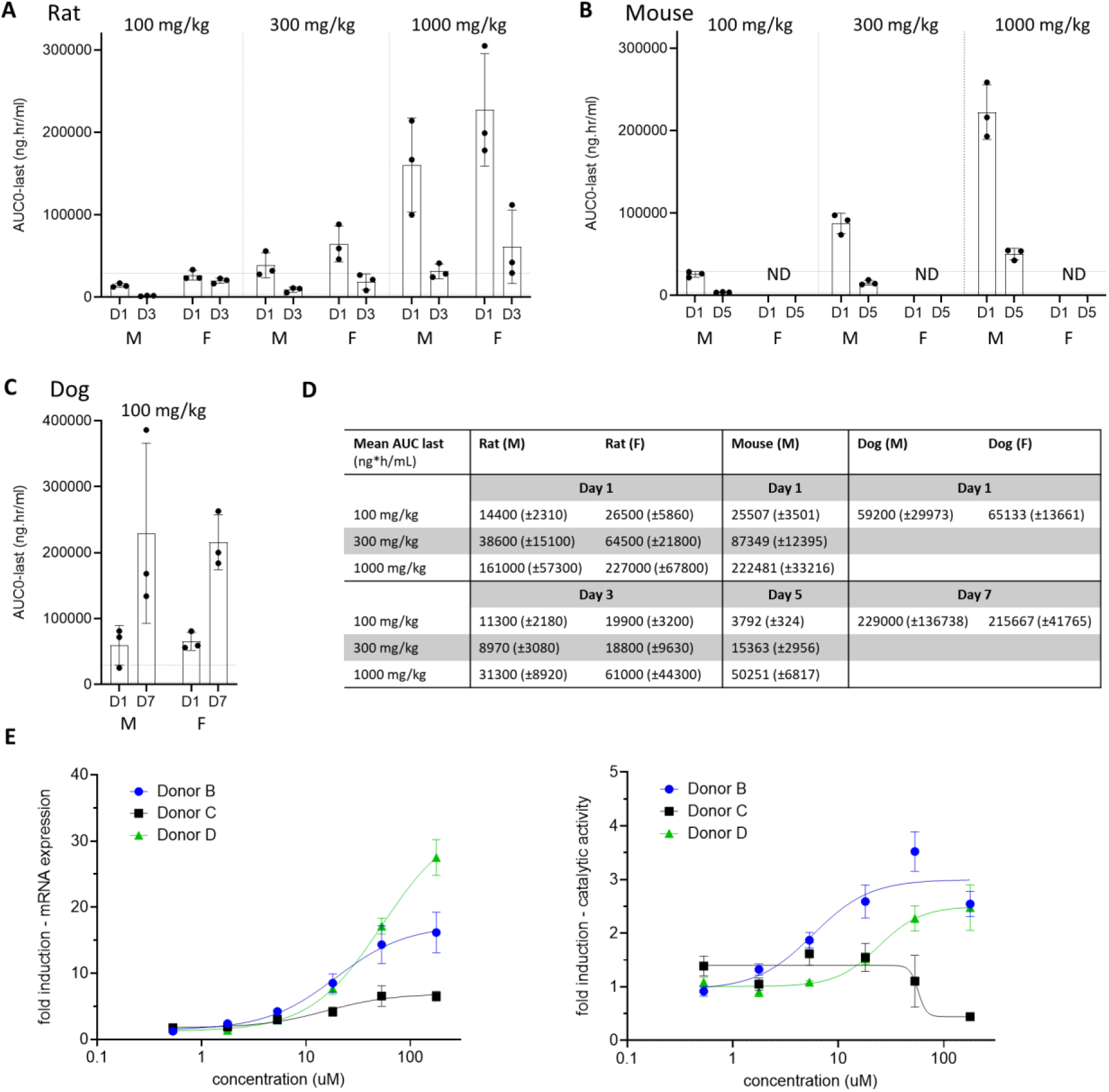
Multi-dosing studies identify rodent-specific decrease in (S)-x38 DNDI-6510 exposures over time. (A) Non-GLP maximum tolerated dose (MTD) study in rat at doses of 100, 300 and 1000 mg/kg showed decreased exposures plotted as AUC_last_ (ng*h/ml) on day 3 in comparison to day 1. The toxicology window (AUC_0-∞, ss_ (µg/mL*h): 3.56 to 28.97) based on the human dose prediction of 125 TID to 1010 mg BID is marked by dotted lines. (B) Multi-day PK study over 5 days in mouse (male only) at 100, 300, and 1000 mg/kg show decreased exposures AUC_last_ (ng*h/ml) on day 5 in comparison to day 1. (C) Multi-day PK study of (S)-x38 DNDI-6510 over 7 days in dog at 100 mg/kg show increased exposures AUC_last_ (ng*h/ml) on day 7 in comparison to day 1. (D) AUC_last_ measurements in MTD studies in rat, and multi-dosing PK studies in mouse (day 1 and day 5) and dog (day 1 and day 7). (E, F) CYP3A4 mRNA fold induction and (F) CYP3A4 catalytic activity in cryopreserved human hepatocytes from three different donors in response to incubation with DNDI-6510.

These results suggested rapid rodent-specific induction *in vivo*. We therefore investigated *in vitro* nuclear hormone receptor (NHR) induction, assessing the impact of (S)-**x38** DNDI-6510 on PXR, CAR, and AhR, across different species. The study demonstrated PXR-linked induction *in vitro*, with 5-fold increases in mouse, 10-fold in rat and dog, and 17-fold increases in human hepatocyte cell lines, respectively (Table S15). Further, (S)-**x38** DNDI-6510 showed a dose-dependent effect on CYP3A4 mRNA expression in cryopreserved hepatocytes from three independent human donors (Figure 6e), and on and CYP3A4 catalytic activity in two of three independent human donors (Figure 6f), whilst no impact on CYP2B6 and CYP1A2 mRNA expression or catalytic activity was observed (Figure S3).

(S)-**x38** DNDI-6510 preclinical development was discontinued based on the inability to exceed the predicted efficacious human exposure in two rodent species upon multi-day dosing.

### Synthesis optimization - MedChem route and solid-state work

The synthetic route to (S)-**x38** DNDI-6510 was optimized and its solid phase behavior explored to identify a stable, non-hydroscopic solid form for exploratory toxicology studies and further testing and development. (S)-**x38** was initially prepared using a long linear sequence with separation of the racemate by chiral chromatography as the final step of the synthesis. While suitable for multi mg amounts of (S)-**x38**, this was deemed too inefficient for larger scale synthesis. The modified chemical route is shown in Figure 7. The key starting materials 2-bromo-4-chlorobenzoic acid, cyclopropyl glycine and bromo isoquinoline were available as articles of commerce. It was found that resolution of the racemate by chiral chromatography could be run earlier in the sequence on the methyl ester analogue of (S)-**x38**, thus avoiding less waste towards the final API structure. This chemistry was used to prepare ∼40 g of enantiopure (S)-**x38**. Unfortunately, while the route shown in Figure 7 was fit for purpose to provide gram amounts of (S)-**x38**, it was too expensive and inefficient to provide 100s grams or kgs for further preclinical development and Phase 1 trials, and was redesigned principally to remove the need for chiral chromatography to separate the enantiomers late on in the synthesis. This development will be described in a later publication.

**Figure 7:**
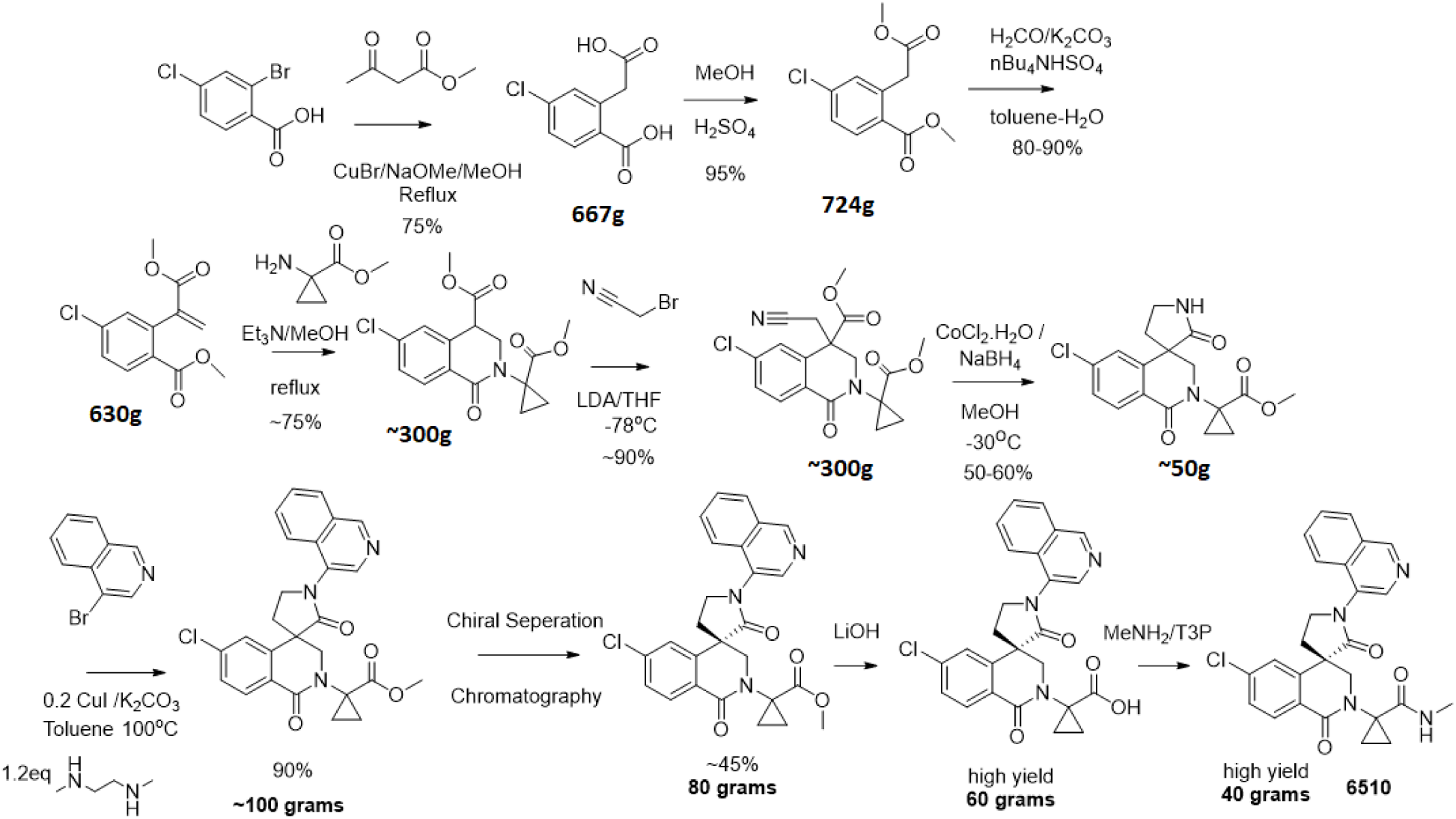
Synthetic route utilized for early (S)-x38 DNDI-6510 batches.

#### Solid state and polymorph screening

The candidate molecule (S)-**x38** was subjected to standard screening protocols to explore solid state behavior and to identify the preferred developable form of the compound. All isolated forms of (S)-**x38** including amorphous form were characterized by appearance, XRPD, TGA, DSC Analysis, Cycling DSC, Microscopy, DVS, ^1^H-NMR, FT-IR, LC purity, and Raman analysis. Extensive studies of a range of counterions and solvents revealed four salts that could be isolated as solids. The sulfate, p-toluene sulfonate, and naphtalene-2-sulfonate salts were obtained as complex mixtures of phases and solvates and were thus discounted. (S)-**x38** was found to form a stable non-solvated, and non-hygroscopic salt at 1:1 stoichiometry salt with HCl, however, it dissociated to monohydrate form (Form 2) in water within 30 minutes. Therefore, HCl salt was not preferred for polymorph screen due to the hydrate formation in a very short time. Polymorph screening using recrystallization and solvent slurry experiments revealed a complex solid form landscape with a total of eleven forms isolated. Most of these were characterized as mixtures/solvates and discarded, except Form 2 being a monohydrate and form 11 being anhydrous pure form.

Short term stability and solubility studies in water, buffer pH 7.4 and simulated fluids (FaSSIF, FeSSIF and FaSSGF) were conducted on Form 2, Form 11, and the amorphous form. All these forms were found stable at 70°C/75% RH and/or 100°C/uncontrolled % RH for 7 days. Solubility of Form 11 and amorphous form is better compared to the poor solubility of Form 2. However, XRPD after 24 hours in dissolution media shows a conversion to Form 2 for all experiments indicating limited stability of Form 11 and amorphous material in aqueous media.

Scale up of Form 11 was investigated using slurry at 50°C and cooling crystallization in acetonitrile; cooling recrystallization with seeding proved to be a reliable and scalable methodology to access pure Form 11. In contrast, production of pure amorphous (S)-**x38** was challenging; only freeze drying or ball milling in the presence of n-heptane (30 min at 30Hz) yielded consistent results.

Therefore, Form 11 was chosen for all future development work as it is the most stable polymorphic form based on the melting temperature (∼271.0°C), improved solubility, favourable stability profile and process scalability. Nevertheless, the conversion of all forms to the highly insoluble (S)-**x38** monohydrate (Form 2) in aqueous media is a concern for future development.

## Discussion

Here, we report the discovery and profiling of a novel SARS-CoV-2 main protease inhibitor (S)-**x38** DNDI-6510. (S)-**x38** addresses the key liabilities of its predecessor, the Moonshot lead (S)-x1, including a positive AMES test as well as a high rodent clearance. (S)-x1 was discovered by the COVID Moonshot consortium based on a structure-based fragment screen and is structurally differentiated to current clinical main protease inhibitors (Figure 1). The SARS-CoV-2 main protease has been exploited as a drug development target by many efforts, with numerous novel SARS-CoV-2 inhibitors in the clinical and preclinical pipeline. The regulatory approved first-generation inhibitors nirmatrelvir/ritonavir (Paxlovid, Pfizer)^3^ and ensitrelvir (Xocova, Shionogi)^4^ have been rapidly followed by second generation inhibitors now in clinical and preclinical development, addressing limitations on antiviral spectrum, pharmacokinetic limitations and resistance profile. Follow-on covalent inhibitors include ibuzatrelvir (PF-0781883)^6^, an optimised nirmatrelvir derivate without the need of ritonavir co-administration, now in Phase III clinical trials^23,24^. Further, a number of preclinical inhibitors that harbor close resemblance to nirmatrelvir have been disclosed^25–29^. Non-peptidomimetic non-covalent inhibitors, falling into the same class as ensitrelvir, include both non-isoquinoline compounds^30–32^, as well as those containing an isoquinoline^13,17,33^. The most recently disclosed non-covalent Mpro series include AVI-4773 and AVI-4516 harbors an isoquinoline moiety^34^, which we also established as a critical moiety for Mpro potency for our scaffold (Figure 2d).

Although the 4-amino-isoquinoline moiety engenders significant potency benefit in our series, we were concerned that, like other amino-heterocycles, it could generate an AMES liability as reported previously^14^. For a community use, oral coronavirus therapeutic, an AMES risk early in discovery would be considered as an unacceptable risk, since key populations would be excluded from using the compound. Indeed, when we tested 4-amino-isoquinoline AMES SAR, including a potent member of our series, we found a positive signal across TA-98 and TA-100 strains (Figure 2b), which was not abrogated by varying the isoquinoline-6 position (Figure 2c). The acyclic central amide linkage is a probable position of metabolism, releasing the amino-isoquinoline that then elicits a positive response in the assay, as detected for **x6** (Figure 2b). Further, despite exploring a wide range of alternative heterocycles, the isoquinoline interactions were found to critically impact enzyme potency of the series (Figure 2d), reflected by multiple other disclosed non-covalent structures harboring an isoquinoline^17,33,34^. A final strategy was locking in the isoquinoline through cyclisation, a commonly used approach to address metabolic liability^35^, but to our knowledge not previously used to address an AMES risk linked to amino-heterocycle release. We had generated a sequence of spirocyclic crystal structures (PDB IDs 7GHU, 7GJC, 7GMJ, 7GK9) that identified the critical carbonyl necessary to maintain binding, by forming the aforementioned hydrogen bond with the backbone NH of Glu166. Importantly, the spirolactam is expected to be metabolically stable; as, even if the amide is cleaved, intramolecular re-cyclisation would be quicker than release of the amino-isoquinoline^36^. Preparation of the spiro-lactam (*S*)-**x29** yielded both high potency and no concerning activity in the AMES test even at high concentrations (Figure 2b). In addition, we were able to develop a synthetic route using an analogue of the spiro-lactam with 4-bromo-isoquinoline which does not generate a risk of 4-amino-isoquinoline contamination in the synthetic intermediates (Figure 7).

We chose to adopt a systematic approach to addressing the different potential sites of metabolism. Multi-factorial optimization of potency and clearance was demonstrated over multiple matched pairs, with spirocyclisation reducing central amide cleavage, deceleration of sidechain hydrolysis through cyclopropanation and blockade of benzylic oxidation. For the side chain hydrolysis, the use of cyclopropyl as a conformationally restricted bio-isosteric replacement is precedented by the work of Wood et al^37^ in their design of Bradykinin B1 Receptor Antagonists. We were not successful in maintaining potency while reducing clearance by attempting to address either isoquinoline N-oxidation or isoquinoline aromatic oxidation. The combined effect of the successful strategies however led to a significant improvement in oral exposure in rodent. Integrating crystallographic screening and structural biology into the medicinal chemistry progression at high scale was crucial for understanding the MPro enzyme SAR and progressing towards optimized leads that were selected for in-depth studies. Further, early rat IV dosing in cassette studies to fast-forward lead optimization was a fruitful strategy to generate a holistic readout and optimize multiple metabolic hotspots in the molecule. Extensive use of matched pair analysis allowed us to quickly zoom in on relevant chances, in particular the introduction of the carbonyl and the cyclisation contributed to the main improvements of metabolic liability, which also appeared to be additive.

Preclinical demonstration of efficacy in an animal mouse model of disease is considered a key step in drug discovery. However, mouse-specific pathways of drug metabolism can impact on exposure, and addressing rapid rodent metabolic clearance is a common problem during lead optimization. Our lead compound (S)-x38 showed excellent *in vitro* profiles in liver microsomes and hepatocytes in higher species including human (Table S8, S9), good pharmacokinetic properties in a second non-rodent species (Table S13), and an acceptable initial human dose prediction (Table S2). Despite significant optimization of (S)-x38 rodent metabolic liabilities, in vivo exposures in wild-type animals were not sufficient to reach sufficient exposures above free antiviral EC90 for >12 hours, allowing for BID dosing (Figure 5b). Based on these observations, we utilized a co-dosing model with the pharmacokinetic enhancer ABT as well as a humanized mouse model (8HUM) as strategies to achieve exposure in mouse for an *in vivo* proof of concept study. Especially the 8HUM model is of translational relevance, as the pharmacokinetics and metabolite profiles were shown to be in much closer alignment with clinical observations than observed for wild-type mice^21^. Indeed, both strategies extended the exposure of (S)-x38 over 3x free antiviral EC90, with the highest tested dose in pharmacokinetic studies for 10 hours in the ABT co-dosing model and for >12 hours in the humanized mouse model, respectively (Figure 5b). However, to achieve a reduction in viral loads *in vivo*, continuous exposure over 3x EC90 was required (Figure 5c). These results are in line with observations reported for nirmatrelvir, where exposures exceeding 3x EC90 were required to achieve *in vitro* efficacy in mice^3^. We conclude that utilizing the 8HUM mouse model to establish in vivo efficacy of a compound with rapid rodent metabolism is a feasible translational strategy.

After thorough characterization of (S)-x38 DNDI-6510, we concluded that despite significant improvements on the series metabolic and genotoxic profile, remaining limitations of the described lead (S)-x38 prevent its translational development in the current environment. Firstly, similarly to ensitrelvir^4^, the antiviral spectrum of (S)-x38 is comparatively narrow; whilst active against all tested SARS-CoV-2 variants as well as SARS-CoV-1, it does not show any broader activity against betacoronavirus MERS and alphacoronavirus 229E in cellular antiviral assays (Figure 4e), offering limited suitability for a pandemic preparedness scenario where a MERS-like coronavirus variant may circulate. Secondly, we observed rapid metabolic induction in MTD studies, with rodent exposures significantly decreased after three to five days of dosing. This decrease in exposure over time prevented achieving sufficient exposures in nonclinical species to consistently explore the targeted human exposures. Based on these considerations, (S)-x38 was not progressed to exploratory toxicology studies. Even though the initial PXR screening result was negative, more rigorous evaluation in a nuclear hormone receptor assay across species suggested that induction may be PXR linked (Table S15). We have also shown that - even though PXR induction was shown in vitro for all species (mouse, rat, dog, and human, Table S15) in a nuclear hormone assay - pronounced *in vivo* induction was only observed in rodent, but not dog (Figure 6a to c). Finally, we demonstrated that (S)-x38 induces CYP3A4 mRNA expression and CYP3A4 catalytic activity in cryopreserved hepatocytes (Figure 6e). Of note, even though it is perceived in the field that CYP3A4 induction occurs over longer time frames, there is published evidence that rapid induction of CYP3A4 enzymatic activity with subsequent impact on drug exposure levels can be trigger in short time frames less than 24 hours^38^.

In line with the open-science spirit of the COVID Moonshot that led to the discovery of (S)-x1, we are disclosing all project data through a Public CDD vault and ChEMBL in line with FAIR principles. These datasets are too comprehensive to disclose in a publication, but provide an open record of multiple other chemistry strategies that were tested during the late lead optimization stages.

We expect this to be a unique resource for computational chemistry education, design and machine learning efforts.

## Materials and methods

### Compound registration and data flow process

All compound designs from the COVID-19 Moonshot medicinal chemistry team, collaborators, and external submitters were captured through the online compound design submission platform (https://covid.postera.ai/covid) as described previously^13^; along with submitter identity, institution, design rationale, and any inspiration fragments. In brief, each submitted batch of related designs received a unique ID including the first three letters of the submitter name and submitter institution, and each compound design submitted received a unique ID (“PostEra ID”) that appended a unique molecule sequence ID within the submission batch ID. Compound designs, synthesized compounds, and compounds with experimental data were tracked with corresponding records in a CDD Vault (Collaborative Drug Discovery Inc.), now available as an publicly accessible vault.

Compounds were initially synthesized and biochemically assayed as racemates, and if active, chirally separated compounds were registered and assayed separately. Absolute stereochemical identity of enantiopure compounds was unknown at time of receipt, therefore, assay data were attached to compound records with specified relative stereochemistry, rather than absolute stereochemistry (e.g., x-ray data for the compound or a close analog). Then, “suspected_SMILES” record was updated along with an articulated rationale in the “why_suspected_SMILES” field.

### Computational methods

#### Synthetic route planning

We use an approach based on the Molecular Transformer technology as described^13^. Our algorithm uses natural language processing to predict the outcomes of chemical reactions and design retrosynthetic routes starting from commercially available building blocks. This proprietary platform was provided by PostEra Inc (https://postera.ai/). Manifold (https://app.postera.ai/manifold/) was built by PostEra Inc. during the COVID Moonshot project to search the entire space of purchasable molecules and find optimal building blocks^13^, and further utilized during this phase of the project.

#### Alchemical free-energy calculations

As described^13^, large-scale alchemical free-energy calculations were conducted in batches (“Sprints”) in which each set of calculations aimed to prioritize compounds that could be produced from a common synthetic intermediate using Enamine’s extensive building block library, resulting in synthetic libraries of hundreds to tens of thousands. Virtual synthetic libraries were organized into a star map, where all transformations were made with respect to a single reference x-ray structure and compound with experimentally measured bioactivity. x-ray structures were prepared using the OpenEye Toolkit SpruceTK with manually controlled protonation states for the key His^41^:Cys^145^ catalytic dyad (variously using zwitterionic or uncharged states) and His^163^ in P1 (which interacts with the 3-aminopyridine or isoquinoline nitrogen in our primary lead series). As the most relevant protonation states were uncertain, when computational resources afforded, calculations were carried out using multiple protonation state variants (His^41^:Cys^145^ either neutral or zwitterionic; His^163^ neutral or protonated) and the most predictive model on available retrospective data for that scaffold selected for nominating prospective predictions for that batch. Initial poses of target compounds were generated via constrained conformer enumeration to identify minimally clashing poses using Omega (from the OpenEye Toolkit) using a strategy that closely follows an exercise described in a blog post by Pat Walters (http://practicalcheminformatics.blogspot.com/2020/03/building-on-fragments-from-diamondxchem_30.html). Alchemical free-energy calculations were then prepared using the open source perses relative alchemical free-energy toolkit^39^ (https://github.com/choderalab/perses), and nonequilibrium switching alchemical free-energy calculations^40^ were run on Folding@home using the OpenMM compute core^41^. Scripts for setting up and analyzing the perses alchemical free-energy calculations on Folding@home, as well as an index of computed datasets and dashboards are available at https://github.com/foldingathome/covid-moonshot Code used for generating the COVID Moonshot alchemical free-energy calculation web dashboards is^21^ available here: https://github.com/choderalab/fah-xchem.

### Chemical methods

#### General compounds synthesis and characterization

All compounds were directly purchased from Enamine Inc., following Enamine’s standard quality control (QC) for compound collections. In addition, in the supplementary chemistry section of the supplementary materials, we discuss the synthesis procedure, as well as liquid chromatography–mass spectrometry (LC-MS) and 1H nuclear magnetic resonance (NMR) characterization of compounds which were discussed in the manuscript with associated bioactivity data.

All COVID Moonshot compounds are publicly available as a screening collection that can be ordered in bulk or as singleton through Enamine. The compound identifiers of the COVID Moonshot collection are in the supplementary data files, together with Enamine’s internal QC data comprising LC-MS spectra for all compounds and NMR spectra for selected compounds.

### Experimental methods

#### Fluorescence Mpro inhibition assay

Compounds were measured in biochemical MPro assays as described previously. In brief, compounds were seeded into assay-ready plates in duplicate (Greiner 384 low volume, cat. no. 784900) using an Echo 555 acoustic dispenser, and dimethyl sulfoxide (DMSO) was back-filled for a uniform concentration in assay plates (DMSO concentration maximum 1%). Dose response assays were performed in 12-point dilutions of twofold, typically beginning at 100 μM. Highly active compounds were repeated in a similar fashion at lower concentrations beginning at 10 μM or 1 μM. Reagents for Mpro assay were dispensed into the assay plate in 10 μl volumes for a final volume of 20 μl.

Final reaction concentrations were 20 mM HEPES pH 7.3, 1.0 mM TCEP, 50 mM NaCl, 0.01% Tween-20, 10% glycerol, 5 nM Mpro, 375 nM fluorogenic peptide substrate ([5-FAM]-AVLQSGFR-[Lys(Dabcyl)]-K-amide). Mpro was pre-incubated for 15 min at room temperature with compound before addition of substrate, and after for 30 minutes. Protease reaction was measured in a BMG Pherastar FS with a 480/520 excitation/emission filter set. Raw data were mapped and normalized to high (Protease with DMSO) and low (No Protease) controls using Genedata Screener software. Normalized data were then uploaded to CDD Vault (Collaborative Drug Discovery). Dose response curves were generated for IC50 using nonlinear regression with the Levenberg–Marquardt algorithm with minimum inhibition = 0% and maximum inhibition = 100%.

#### High-throughput X-ray crystallography

High-throughput X-ray crystallography for this project was performed as described previously^13^. In brief, purified protein at 24 mg/ml in 20 mM HEPES pH 7.5, 50 mM NaCl buffer was diluted to 12 mg/ml with 20 mM HEPES pH 7.5, 50 mM NaCl before performing crystallization using the sitting-drop vapor diffusion method with a reservoir solution containing 11% PEG 4 K, 5% DMSO, 0.1 M MES pH 6.5. Crystals of Mpro in the monoclinic crystal form (C2), with a single monomer in the asymmetric unit, were grown with drop ratios of 0.15 μl protein, 0.3 μl reservoir solution, and 0.05 μl seeds prepared from previously produced crystals of the same crystal form. Crystals in the orthorhombic crystal form (P212121), with the Mpro dimer present in the asymmetric unit, were grown with drop ratios of 0.15 μl protein, 0.15 μl reservoir solution, and 0.05 μl seeds prepared from crystals of an immature Mpro mutant in the same crystal form^42^.

Compounds were soaked into crystals by adding compound stock solutions directly to the crystallization drops using an ECHO liquid handler. In brief, 40 to 90 nl of DMSO solutions (between 20 and 100 mM) were transferred directly to crystallization drops giving a final compound concentration of 2 to 20 mM and DMSO concentration of 10 to 20%. Drops were incubated at room temperature for ∼1 to 3 hours before mounting and flash cooling in liquid nitrogen without the addition of further cryoprotectant. Data were collected at Diamond Light Source on the beamline I04-1 at 100 K and processed with the fully automated pipelines at Diamond^43,44^. Further analysis was performed using XChemExplorer^45^ with electron density maps generated using DIMPLE (http://ccp4.github.io/dimple/). Ligand-binding events were identified using PanDDA^46^ (https://github.com/ConorFWild/pandda), and ligands were manually modeled into PanDDA-calculated event maps or electron density maps using Coot^47^. Ligand restraints were calculated with ACEDRG ^48^ or GRADE [grade v. 1.2.19 (Global Phasing Ltd., Cambridge, UK, 2010)] and structures refined with Buster [Buster v. 2.10.13 (Cambridge, UK, 2017)]. Models and quality annotations were reviewed using XChemReview, Buster-Report [Buster v. 2.10.13 (Cambridge, UK, 2017)] and Mogul as described previously^13^. Coordinates, structure factors and PanDDA event maps for all datasets are available from the Protein Data Bank.

#### Cytopathic effect assay, hACE2-TMPRSS2 cells (Katholieke Universiteit Leuven)

##### Virus isolation and virus stocks

All virus-related work was conducted in the high-containment BSL3 facilities of the KU Leuven Rega Institute (3CAPS) under licenses AMV 30112018 SBB 219 2018 0892 and AMV 23102017 SBB 219 2017 0589 according to institutional guidelines.

Viral strains were described previously^49^. The SARS-CoV-2 GHB strain (GHB-03021/2020, GISAID: EPI_ISL_407976 | 2020-02-03) was recovered from a nasopharyngeal swab taken from an asymptomatic patient returning from Wuhan, China. Virus stocks were inoculated on HuH-7 cells and then passaged seven times on VeroE6–eGFP cells. SARS-CoV-2 strains belonging to the alpha, beta, delta and omicron variants of concern were recovered from nasopharyngeal swabs of human cases confirmed by quantitative PCR with reverse transcription (RT–qPCR). Virus stocks were generated by passaging the virus on Calu-3 cells followed by production of a screening virus stock on A549ACE2+TMPRSS2 cells. Median tissue culture infectious doses (TCID50) was defined by end-point titration. The sequences of the passage 0 of these strains are available through GISAID: Alpha/B.1.1.7 (hCoV19/Belgium/rega-12211513/2020; EPI_ISL_791333), beta/B.1.351 (hCoV19/Belgium/rega-1920/2021; EPI_ISL_896474); Delta/B.1.617.2 (EPI_ISL_2425097); Omicron BA.5 (EPI_ISL_14782497); and XBB1.5 (EPI_ISL_17273054). The sequences of BA.2.86 (SARS-CoV-2/hu/DK/SSI-H135) and EG.5.1 (SARS-CoV-2/hu/DK/SSI-H121) variants are available in the European Nucleotide Archive under the project number PRJEB67449 with accession numbers OY747653 and OY747654, respectively. All used cell lines have been described previously^49^.

##### SARS-CoV-2 A549-ACE2-TMPRSS2 assay

A549-Dual hACE2-TMPRSS2 cells obtained by Invitrogen (cat. no. a549d-cov2r) were cultured in DMEM 10% FCS (Hyclone) supplemented with 10 μg/ml blasticidin (Invivogen, ant-bl-05), 100 μg/ml hygromycin (Invivogen, ant-hg-1), 0.5 μg/ml puromycin (Invivogen, ant-pr-1) and 100 μg/ml zeocin (Invivogen, ant-zn-05). For antiviral assay, cells were seeded in assay medium (DMEM 2%) at a density of 15,000 cells per well. One day after, compounds were serially diluted in assay medium (DMEM supplemented with 2% v/v FCS) and cells were infected with their respective SARS-CoV-2 strain at a MOI of ∼0.003 TCID50/ml. On day 4 pi., differences in cell viability caused by virus-induced CPE or by compound-specific side effects were analyzed using MTS as described previously^50^. Cytotoxic effects caused by compound treatment alone were monitored in parallel plates containing mock-infected cells.

#### SARS-CoV-2 in vivo antiviral efficacy (Mount Sinai)

##### In vivo animal efficacy

All in vivo antiviral studies were performed as described previously^51^. In brief, experiments were executed in the animal biosafety level 3 (BSL3) facility at the Icahn school of Medicine in Mount Sinai Hospital, New York City under protocols approved by the Institutional Animal Care and Use Committee (IACUC). Two studies to evaluate the in vivo efficacy of (S)-x38 as antiviral in SARS-CoV-2 infection were performed: (i) in a wild-type 129/S mouse model (the Jackson laboratory strain #002448), in which (S)-x38 was co-dosed with an ABT inhibitor. Second, (S)-x38 was dosed in a humanized mouse model described previously^21^ and purchased through Phaser Ltd.

1. For the first experiment, 45 male and female 12-week-old specific pathogen–free 129/S mice were utilized. (S)-x38 was administrated orally (p.o.) by gavage, at doses of 15, 50, and 150 mg/kg twice a day for 3 days, with co-dosing with 1-aminobenzotriazole (ABT) co-dosed at 50 mg/kg with the morning dose. A vehicle of 1% Methylcellulose, 0.1% Tween 80 was used.
2. For the second animal study, we utilized 45 male and female humanized mice (8HUM) on a C68Bl6 background supplied by Phaser Ltd. (S)-x38 was administrated orally (p.o.) by gavage, at doses of 50, 150, and 300 mg/kg twice a day for 3 days. A vehicle of 0.5% Methylcellulose, 0.05% Tween 80 was used.

For both groups, we administered vehicle, (S)-x38 or positive control nirmatrelvir at 600 mg/kg, 2 hours before intranasal infection with 2.5 × 10^4^ PFU of mouse-adapted SARS-CoV-2 in 50 μl of PBS. 2 uninfected mice were included per group were included for satellite measurements, that were cheek bled once a day prior to the morning dose to achieve trough pharmacokinetic measurements. Mice were anesthetized with a mixture of ketamine/xylazine before each intranasal infection. Each day mouse bodyweight was measured. 3 days post infection (dpi) animals were humanely euthanized.

##### Lung viral titer analysis

Whole left lungs were harvested and homogenized in PBS with silica glass beads then frozen at −80°C for viral titration via TCID50. Briefly, infectious supernatants were collected at 48h post infection and frozen at −80°C until later use. Infectious titers were quantified by limiting dilution titration using Vero E6/TMPRSS2 cells. Briefly, Vero E6 cells were seeded in 96-well plates at 20,000 cells/well. The next day, SARS-CoV-2-containing supernatant was applied at serial 10-fold dilutions ranging from 10^−1^ to 10^−6^ and, after 3 days, viral cytopathic effect (CPE) was detected by staining cell monolayers with crystal violet. TCID50/ml were calculated using the method of Reed and Muench. The Prism software (GraphPad) was used to determine differences in lung titers using an unpaired T test on log-transformed data.

##### Histopathology

Paraffin-embedded lung tissue blocks for mouse lungs were cut into 5μm sections. Sections were stained with hematoxylin and eosin (H&E) and analyzed by Histowiz (Brooklyn, NY). Digital light microscopic scans of whole lung processed in toto were examined by an experienced veterinary pathologist. Hematoxylin Eosin stained sections of lung from 8HUM mice were examined by implementing a semi quantitative, 5-point grading scheme (0 - within normal limits, 1 - mild, 2 - moderate, 3 - marked, 4 - severe); considering four different histopathological parameters: 1) perivascular inflammation 2) bronchial or bronchiolar epithelial degeneration or necrosis 3) bronchial or bronchiolar inflammation and 4) alveolar inflammation. These changes were absent (grade 0) in lungs from vehicle and plitidepsin treated uninfected mice from groups that were utilized for this assessment. Data was analyzed by two-way ANOVA (*P < 0.05, **P < 0.01, ***P < 0.001, and ****P < 0.0001).4. Preclinical profiling

#### Plasma protein binding

Tests were conducted by TCG Life Sciences, India, using the equilibrium dialysis method and a 96-well plate format. The test compounds (at 1 μM, in duplicate) in solution in plasma (Balb/c mouse,, Sprague-Dawley rat, Beagle dog, monkey, minipig or human) were dialyzed against PBS (pH 7.4) on a rotating plate incubated for 6 h at 37 °C in an Equilibrium dialysis block: Pierce (Cat No: 89811); Dialysis chamber: Pierce (Cat No:89809) alongside ranitidine, propranolol and warfarin as QC standards. Following the precipitation of protein with ice-cold acetonitrile, samples were mixed and then centrifuged at 4000 rpm for 20 minutes at 15°C. Subsequently, the amount of compound present in each compartment was quantified by LC-MS/MS.

#### Protease selectivity panel

Compounds were tested at Nanosyn, USA, in 12-point concentration-response format against 13 human proteases. Test compounds were diluted in 100% DMSO using 3-fold dilution steps, with final compound concentrations ranging from 100 µM to 0.565 nM. Compounds were tested in a single well for each dilution, and the final concentration of DMSO in all assays was kept at 1%.

For proteases Calpain1, Sigma; CathepsinB, R&D Biosystems; Cathepsin K, L and S, Calbiochem, enzyme concentrations of 3.6, 12.5, 1.5, 0.67 and 10 nM for 1-3 hours, respectively. Reference compound, E-64, was tested in an identical manner. IC50 curves were generated for all tested proteases.

#### Metabolic stability in hepatic microsomes

Tests were performed by TCG Life Sciences, India. Test compounds (at 1 μM) in duplicate or positive controls (atenolol, propanolol, diclofenac, and verapamil) were incubated at 37 °C with liver microsomes from Balb/C mouse, Sprague-Dawley rat, Beagle dog, monkey or human in the presence of a NADPH regenerating system (NRS) and phosphate buffer (100 mM, pH 7.4) at 0.4 mg/mL microsomal protein. The samples were removed at time intervals from 0, 5, 10, 20, 30 and 60 minutes and immediately mixed with cold methanol supplemented or not with acid (3% formic acid) and centrifuged prior to analysis by LC-MS/MS using propranolol as internal standards.

#### Metabolic stability in hepatic microsomes

Metabolic stability in liver microsomes derived from humanized mice (8HUM were performed as described previously^21^. In brief, test compound (S)-x38 at 0.5 µM final concentration was combined with microsomes in buffer (0.5 mg/mL 50 mM potassium phosphate, pH 7.4) and the reaction initiated with addition of excess NADPH (final concentration 0.8 mg/mL). Aliquots of 50 µL were removed from the incubation mixture at 0, 3, 6, 9, 15, and 30 min, mixed with acetonitrile (100 µL), and kept on ice. After all samples were collected, 250 µL of 20% acetonitrile was added to each and the analysis plate was centrifuged for 10 min at 3,000 × g at ambient temperature and analyzed immediately.

#### Metabolic stability in hepatocytes

Tests were performed by Cyprotex Discovery Ltd. (UK). Following a viability check of cryopreserved hepatocytes from Sprague-Dawley rat, Beagle dog, or human, test compounds (at 1 μM) or positive controls 7-ethoxycoumarin and 7-hydroxycoumarin (at 3 μM) were incubated at 37 °C, 5% CO2 with hepatocytes in Williams’ Medium E. Following incubation of 15, 30, 60, 90, 120, 180, and 240 min, reactions were stopped by the addition of acetonitrile, the samples were centrifuged, and LC-MS/MS analysis was conducted using tolbutamide as an internal standard.

#### Metabolite identification in microsomes

Metabolite profiling in liver microsomes has been conducted at Cyprotex Discovery Ltd. (UK) on the test compounds in rat microsomes. The 45-minute time point was compared against the 0-minute control sample to establish which and how many metabolites were formed. The metabolites found have been displayed in this report as extracted ion chromatograms (XIC) and representative low and high energy mass spectra. The areas and percentages reported for the parent and metabolites have been calculated using the XIC data; it has been assumed that each metabolite has the same ionisation efficiency and that the sensitivity of the metabolite has not been affected by the biotransformation. Any potential matrix effects have also been assumed to be consistent with the parent molecule. Metabolites observed at greater than 1% of the total of drug related material are reported. When observed in other species, metabolites observed at less than 1% were also reported. Three metabolites have been found and reported in rat microsomes.

#### Metabolite identification in hepatocytes

Test compounds were tested at Novartis (in-kind contribution) for metabolic stability using cryopreserved hepatocytes from mouse, rat, dog, minipig, monkey and human. 1 µM test article was incubated with 1 Mio cells/mL (L-15 medium) at 37°C in 96 plates and shaking with 1000 rpm for 1, 10, 20, 40, 60 and 80 min. Analysis was by LC-MS doing full-scan with data-dependent MSMS, with LC separation using a generic gradient up to 10 min using a Q-Exactive. Data were analyzed using a SSID approach, and Mass-MetaSite used to identify metabolites and manual evaluation for cross species comparison.

#### AMES testing

AMES testing was performed at Cyprotex Ltd. Approximately ten million bacteria were exposed in triplicate to the test agent (six concentrations), a negative control (vehicle) and a positive control for 90 minutes in medium containing a low concentration of histidine (sufficient for about 2 doublings.) The cultures were then diluted into indicator medium lacking histidine, and dispensed into 48 wells of 384-well plates (micro-plate format, MPF). The plates were incubated for 48 hrs at 37°C, and cells that underwent a reversion grew, resulting in a color change in the wells with growth. The number of wells showing growth were counted and compared to the vehicle control. An increase in the number of colonies of at least two-fold over baseline (mean + SD of the vehicle control) and a dose response indicate a positive response. An unpaired, one-sided Student’s T-test was used to identify the conditions that were significantly different from the vehicle control. Where indicated, S9 fraction from the livers of Aroclor 1254-treated rats were included in the incubation at a final concentration of 4.5%. A NADPH-regenerating system was also included to ensure a steady supply of reducing equivalents. Strains used in this study: S. typhimurium TA98: hisD3052, rfa, uvrB / pKM101; detects frame-shift mutations.S. typhimurium TA100: hisG45, rfa, uvrB / pKM101; detects base-pair substitutions.

#### Nuclear Hormone receptor profiling

Nuclear hormone receptors PXR and AhR were profiled at Cyprotex Discovery Ltd. (UK) across species (human, dog, mouse, rat), whilst CAR receptors were profiled species-specific, for human (CAR1, CAR3); dog (CAR3); mouse (CAR1) and rat (CAR3). Species-specific AXR, CAR and PXR cells were seeded in 96-well plates and recovered overnight at 37 °C in 5 % CO2 for 24 hr prior to dosing. Cells were dosed with increasing concentrations of compound (concentrations of 0.4, 1, 4, 10, 40 and 100 μM) with 2 replicates per concentration. Positive controls in duplicate included Rifampicin for human and dog PXR (concentrations of 0.025, 0.1, 0.25, 1, 2.5, 10 and 25 μM), Pregnenolone-16-α-carbonitrile (PCN) for mouse and rat PXR (concentrations of 0.01, 0.05, 0.1, 0.5, 1 and 5 μM), 3-Methylcholanthrene (3-MC) for human, mouse, dog, and rat AhR (concentrations of 0.01, 0.05, 1, 5, 10 and 20 μM), Phenytoin for human CAR1 (0.005, 0.1, 0.5, 1, 5, 10, 20 and 100 μM), 6-(4-Chlorophenyl)imidazo[2,1-b][1,3]thiazole-5-carbaldehyde O-(3,4-dichlorobenzyl)oxime (CITCO) for human, rat and dog CAR3 (compound concentrations of 0.005, 0.1, 0.5, 1, 5, 10 and 20 μM),3,3′,5,5′-Tetrachloro-1,4-bis(pyridyloxy)benzene (TCPOBOP) for mouse CAR1 (concentrations of 0.01, 0.1, 0.5, 1, 5, 10 and 20 μM), and vehicle control, respectively. Compounds and cells were incubated at 37 °C for 48 hr. Following incubation, culture media was discarded and cell viability assessed fluorimetrically using CellTiter-FluorTM, and luciferase activity is measured using ONE-GloTM. A fold increase in transcriptional activation of PXR above solvent control was calculated for two replicates for each measured concentration (0.4, 1, 4, 10, 40 and 100 uM), along with derived EC50 value determined where appropriate. Data were expressed as fold activation relative to the vehicle control, following normalisation of luciferase activity for cell number.

#### CYP inhibition

The study was conducted by Cyprotex Discovery Ltd. (UK), using the time-dependent (TDI) method. To assess the potential reversible and time dependent inhibition of cytochrome P450 isoforms using isoform-specific probe substrates, six test article concentrations (typically 0.1, 0.25, 1, 2.5, 10, 25 µM; final DMSO concentration 0.25 %), 0.1 M phosphate buffer pH 7.4 and human liver microsomes were either pre-incubated for 30 min in the absence and presence of NADPH or undergo a 0 min pre-incubation. At the end of the pre-incubation period, the appropriate isoform-specific probe substrate (CYP1A2 Phenacetin / Acetaminophen, CYP2B6 Bupropion / Hydroxybupropion; CYP2C8 Amodiaquine / N-Desethylamodiaquine; CYP2C9 Diclofenac / 4-Hydroxydiclofenac, CYP2C19 S-Mephenytoin / 4-Hydroxymephenytoin; CYP2D6 Dextromethorphan / Dextrorphan, CYP3A4 Midazolam / 1-Hydroxymidazolam and Testosterone / 6ß-Hydroxytestosterone) and NADPH (1 mM) are then added and the samples incubated at 37 °C for the required incubation time. Positive control time dependent inhibitors for each CYP isoform are screened alongside the test article (CYP1A2, Furafylline; CYP2B6, ThioTEPA; CYP2C8, Gemfibrozil 1-*O*-β-glucuronide; CYP2C9, Tienilic acid; CYP2C19 Fluoxetine; CYP2D6, Paroxetine; CYP3A4; Verapamil). The reactions were terminated by methanol, and plates centrifuged at 2500 rpm for 30 min at 4°C and an aliquot of the supernatant is transferred to fresh 96-well plates. Formic acid in deionised water (final concentration 0.1%) was added to the supernatants prior to analysis of isoform specific metabolites by LC-MS/MS. A decrease in the formation of the metabolite compared to the vehicle control is used to calculate an IC50 value (test article concentration which produces 50% inhibition) for each experimental condition

### In vivo studies

#### PK studies in mice

Pharmacokinetic experiments in humanized mice (8HUM) were performed at Dundee University as described previously^21^. In brief, 8HUM breeding colony were maintained at Charles River Laboratories, and all pharmacokinetic experiments carried out at the University of Dundee under the Animals (Scientific Procedures Act) 1986, as amended in 2012, and in accordance with the European Union Directive (2010/63/EU). All studies were approved by the local Ethical Review Committee and undertaken in line with the 3Rs principles of replacement, reduction, and refinement (https://www.nc3rs.org.uk). Female mice were used in all experiments. Dose levels of 50 and 150 mg/kg were administered by gavage, and serial blood samples (10 µL) were taken from the tail vein at the timepoints shown, diluted into 9 volumes of Milli-Q water, and stored at −20 °C prior to bioanalysis. Analysis was carried out by LC-MS/MS, and non-compartmental analysis, modeling, and simulation of PK was carried out using Phoenix WinNonlin version 8.3.1.5014 (Certara) as described previously^21^.

#### PK studies in rat

Single dose rat PK studies were performed at Aptuit, Italy and TCG Life Sciences, India.

At Aptuit, (S)-x38 was dosed to male Sprague Dawley rats (n=3 per group). IV dosing was at 1 mg/kg (20% w/w propylene glycol (PG), 10% w/w Cremophor RH40 and 70% w/w sterile water for injection, 0.2 mg/mL), with serial sampling performed at 0.083, 0.25, 0.5, 1, 3, 6, 8 and 24 hours after dosing; and PO dosing at 2, 5, and 10 mg/kg (2.5% w/v Vit E TPGS in sterile water for injection, at 0.2, 0.5, 1 mg/mL) with 0.25, 0.5, 1, 2, 4, 6, 8 and 24 hours after dosing. PK samples were collected into Potassium EDTA tubes, placed on wet ice immediately and stored at −80 °C until bioanalysis using LC-MS/MS. PK parameters were calculated using the Phoenix WinNonlin software.

At TCG Life Sciences, (S)-x38 was dosed to male Sprague Dawley rats (n=3 per group) orally at 30 and 60 mg/kg (1 % HPMC in water with 1 % tween 80, pH 5-5.5) with sampling performed at 0, 0.083, 0.25, 0.5, 1, 3, 6, 8 and 24 hours after dosing. For 10 mg/kg Bioanalysis was performed using LC-MS/MS, with PK parameters calculated using the Phoenix WinNonlin software version 8.1.

#### PK studies in dog

Single dose dog PK studies were performed at Aptuit, Italy. (S)-**x38** DNDI-6510 was administered to male beagle dogs (n=3 per group, study number VNG12816), IV at 1 mg/kg (5% w/w DMSO and 95% w/w saline, 0.4 mg/mL) with serial sampling at 0.083, 0.25, 0.5, 1.0, 3.0, 6.0, 8.0 and 24 hours after dosing; and PO at 2, 5, and 10 mg/kg (2.5% w/v Vit E TPGS in sterile water for injection, 0.4, 1 and 2 mg/mL) with serial sampling performed at 0.25, 0.5, 1.0, 2.0, 4.0, 6.0, 8.0 and 24 hours after dosing. A cross-over design was used, with a maximum of 3 dose increments and a wash-out period of at least 6 days before each new dose. PK samples were collected into Potassium EDTA tubes, placed on wet ice immediately and stored at −80 °C until bioanalysis using LC-MS/MS. PK parameters were calculated using the Phoenix WinNonlin software.

### Human PK predictions

The human dose was estimated for: once-daily administration, with the requirement to achieve a free steady state plasma concentration at the nadir (C_min_) equal to the 3-fold of the in vitro EC_90_ in the antiviral SARS-CoV-2 A549 cell assay. Additional predictions were run for 1-fold of the in vitro EC_90_. The prediction of human Cl and Vss was based on methods reported by Lombardo *et al*.,^52,53^. The prediction was calculated using in vitro plasma protein binding data across species and 10% FCS (fetal calf serum, to calculate free drug in the EC_90_ assay), as well as rat and dog PK data.

## Author contributions

Conceived the idea for the project: A.V.D, E.G, B.P, P.S; Performed the chemical designs and synthesis strategy: E.G. M.R, A.A.L, R.P.R, B.A.L,; Conducted the FEP computational analysis: J.C; Developed or designed the methodology and assays/models used in these studies: A.V.D., P.S., E.G., F.B.C., S.B., P.G., R.K., A.W., P.R., A.S.; performed the in vivo efficacy studies: B.L.M, K.W; Performed the in vitro cellular efficacy assays: L.V, M.L., D.J.,; Performed the in vitro biochemical assays: S.D., N.L., H.B; Performed the structural biology assays and analysis: D.F, L.K., B.H.B; Performed the in vitro and in vivo ADME and PK experiments: A.K.M., K.D.R; Conducted the human PK and dose predictions: G.F, Performed the visualization of the data: A.V.D, E.G, P.S; Data release work: T.S., J.C., A.A.L., M.R.,; Responsible for the administration of the project: A.V.D, P.S., P.P., P.R, S.B., A.S, A.W., P.G., R.K.; Supervised the studies: F.V.D, J.N, H.B., N.L., J.C, K.D.R; Involved in funding acquisition: A.V.D, B.P., P.S, J.C., A.A.L.; Wrote the original draft: E.G, A.V.D, P.S.; Reviewed and edited the original draft: D.F., M.R., E.G., A.V.D., P.S.; All authors read, revised, and authorized the manuscript before submission.

## Acknowledgements

The COVID Moonshot project is particularly grateful to UCB Pharma Ltd. and UCB SA for the support from their Medicinal and Computational Chemistry groups, to the Novartis Institute for Biomedical Research for generous in-kind ADME and PK contributions, to Takeda for in-kind contribution of antiviral assays/pan-corona biochemical assays, and to Nanosyn for protease panel assays. We thank CDD Vault and OpenEye Scientific for their in-kind contribution allowing the consortium to use their software. We also thank the numerous volunteers that contributed compound designs to the COVID Moonshot, the citizen scientist volunteers of Folding@home for donating their computing resources, and Amazon Web Services for key support of Folding@home infrastructure. We thank Louise Burrows (DNDi) for critical advice on the draft manuscript.

## References

1. COVID-19 deaths | WHO COVID-19 dashboard. Datadot https://data.who.int/dashboards/covid19/deaths.

2. Al-Aly, Z. et al. Long COVID science, research and policy. Nat Med 30, 2148–2164 (2024).

3. Owen, D. R. et al. An oral SARS-CoV-2 Mpro inhibitor clinical candidate for the treatment of COVID-19. Science 374, 1586–1593 (2021).

4. Unoh, Y. et al. Discovery of S-217622, a Noncovalent Oral SARS-CoV-2 3CL Protease Inhibitor Clinical Candidate for Treating COVID-19. J. Med. Chem. 65, 6499–6512 (2022).

5. Pepperrell, T., Ellis, L., Wang, J. & Hill, A. Barriers to Worldwide Access for Paxlovid, a New Treatment for COVID-19. Open Forum Infect Dis 9, ofac174 (2022).

6. Allerton, C. M. N. et al. A Second-Generation Oral SARS-CoV-2 Main Protease Inhibitor Clinical Candidate for the Treatment of COVID-19. J. Med. Chem. 67, 13550–13571 (2024).

7. Chan, P. L. S. et al. Dosing recommendation of nirmatrelvir/ritonavir using an integrated population pharmacokinetic analysis. CPT Pharmacometrics Syst Pharmacol 12, 1897–1910 (2023).

8. Shionogi Receives Award Through BARDA’s Rapid Response Partnership Vehicle to Advance Long-Acting Formulation of S-892216, an Antiviral for COVID-19 Pre-Exposure Prophylaxis in At-Risk Populations. https://www.shionogi.com/us/en/news/2025/01/shionogi-receives-award-through-bardas-rapid-response-partnership-vehicle-to-advance-long-acting-formulation-of-s-892216-an-antiviral-for-covid-19-pre-exposure-prophylaxis-in-at-risk-populations.html.

9. Shionogi Announces Global Phase 3 Trial Demonstrates Post-Exposure Prophylactic Use of Ensitrelvir Prevents Symptomatic COVID-19. https://www.shionogi.com/global/en/news/2024/10/20241029.html.

10. Bouzidi, H. S. et al. Generation and evaluation of protease inhibitor-resistant SARS-CoV-2 strains. Antiviral Research 222, 105814 (2024).

11. Krismer, L. et al. Study of key residues in MERS-CoV and SARS-CoV-2 main proteases for resistance against clinically applied inhibitors nirmatrelvir and ensitrelvir. Npj Viruses 2, 23 (2024).

12. Garcia-Blanco, M. A., Ooi, E. E. & Sessions, O. M. RNA Viruses, Pandemics and Anticipatory Preparedness. Viruses 14, 2176 (2022).

13. Boby, M. L. et al. Open science discovery of potent noncovalent SARS-CoV-2 main protease inhibitors. Science 382, eabo7201 (2023).

14. Tichenor, M. S. et al. Heteroaryl urea inhibitors of fatty acid amide hydrolase: Structure– mutagenicity relationships for arylamine metabolites. Bioorganic & Medicinal Chemistry Letters 22, 7357–7362 (2012).

15. Birch, A. M. et al. Rationally designing safer anilines: the challenging case of 4-aminobiphenyls. J Med Chem 55, 3923–3933 (2012).

16. Shamovsky, I. et al. Explanation for main features of structure-genotoxicity relationships of aromatic amines by theoretical studies of their activation pathways in CYP1A2. J Am Chem Soc 133, 16168–16185 (2011).

17. Luttens, A. et al. Ultralarge Virtual Screening Identifies SARS-CoV-2 Main Protease Inhibitors with Broad-Spectrum Activity against Coronaviruses. J Am Chem Soc 144, 2905–2920 (2022).

18. Bruno, I. J. et al. New software for searching the Cambridge Structural Database and visualizing crystal structures. Acta Crystallogr B 58, 389–397 (2002).

19. von Delft, F. et al. A white-knuckle ride of open COVID drug discovery. Nature 594, 330–332 (2021).

20. Douangamath, A. et al. Crystallographic and electrophilic fragment screening of the SARS-CoV-2 main protease. Nat Commun 11, 5047 (2020).

21. MacLeod, A. K. et al. Acceleration of infectious disease drug discovery and development using a humanized model of drug metabolism. Proceedings of the National Academy of Sciences 121, e2315069121 (2024).

22. Balani, S. K. et al. EFFECTIVE DOSING REGIMEN OF 1-AMINOBENZOTRIAZOLE FOR INHIBITION OF ANTIPYRINE CLEARANCE IN GUINEA PIGS AND MICE USING SERIAL SAMPLING. Drug Metabolism and Disposition 32, 1092–1095 (2004).

23. Mortezavi, M. et al. Virologic Response and Safety of Ibuzatrelvir, A Novel SARS-CoV-2 Antiviral, in Adults With COVID-19. Clin Infect Dis 80, 673–680 (2025).

24. Pfizer. AN INTERVENTIONAL EFFICACY AND SAFETY, PHASE 3, DOUBLE-BLIND, 2-ARM STUDY TO INVESTIGATE ORALLY ADMINISTERED IBUZATRELVIR COMPARED WITH PLACEBO IN NON-HOSPITALIZED SYMPTOMATIC ADULT AND ADOLESCENT PARTICIPANTS WITH COVID-19 WHO ARE AT HIGH RISK OF PROGRESSING TO SEVERE ILLNESS. https://clinicaltrials.gov/study/NCT06679140 (2025).

25. Dayan Elshan, N. G. R., et al. Discovery of CMX990: A Potent SARS-CoV-2 3CL Protease Inhibitor Bearing a Novel Warhead. J Med Chem 67, 2369–2378 (2024).

26. Westberg, M. et al. An orally bioavailable SARS-CoV-2 main protease inhibitor exhibits improved affinity and reduced sensitivity to mutations. Sci Transl Med 16, eadi0979 (2024).

27. Huang, C. et al. A new generation Mpro inhibitor with potent activity against SARS-CoV-2 Omicron variants. Signal Transduct Target Ther 8, 128 (2023).

28. Shurtleff, V. W. et al. Invention of MK-7845, a SARS-CoV-2 3CL Protease Inhibitor Employing a Novel Difluorinated Glutamine Mimic. J Med Chem 67, 3935–3958 (2024).

29. Sun, J. et al. A novel, covalent broad-spectrum inhibitor targeting human coronavirus Mpro. Nat Commun 16, 4546 (2025).

30. Papini, C. et al. Proof-of-concept studies with a computationally designed Mpro inhibitor as a synergistic combination regimen alternative to Paxlovid. Proceedings of the National Academy of Sciences 121, e2320713121 (2024).

31. Kneller, D. W. et al. Structural, Electronic, and Electrostatic Determinants for Inhibitor Binding to Subsites S1 and S2 in SARS-CoV-2 Main Protease. J. Med. Chem. 64, 17366–17383 (2021).

32. Jacobs, L. et al. Design and Optimization of Novel Competitive, Non-peptidic, SARS-CoV-2 Mpro Inhibitors. ACS Med Chem Lett 14, 1434–1440 (2023).

33. Hou, N. et al. Development of Highly Potent Noncovalent Inhibitors of SARS-CoV-2 3CLpro. ACS Cent Sci 9, 217–227 (2023).

34. Detomasi, T. C. et al. Structure-based discovery of highly bioavailable, covalent, broad-spectrum coronavirus MPro inhibitors with potent in vivo efficacy. Sci Adv 11, eadt7836 (2025).

35. Tang, K., Wang, S., Gao, W., Song, Y. & Yu, B. Harnessing the cyclization strategy for new drug discovery. Acta Pharmaceutica Sinica B 12, 4309–4326 (2022).

36. Imming, P., Klar, B. & Dix, D. Hydrolytic Stability versus Ring Size in Lactams: Implications for the Development of Lactam Antibiotics and Other Serine Protease Inhibitors. J. Med. Chem. 43, 4328–4331 (2000).

37. Wood, M. R. et al. Cyclopropylamino acid amide as a pharmacophoric replacement for 2,3-diaminopyridine. Application to the design of novel bradykinin B1 receptor antagonists. J Med Chem 49, 1231–1234 (2006).

38. Iwanaga, K., Honjo, T., Miyazaki, M. & Kakemi, M. Time-dependent changes in hepatic and intestinal induction of cytochrome P450 3A after administration of dexamethasone to rats. Xenobiotica 43, 765–773 (2013).

39. Rufa, D. A. et al. Towards chemical accuracy for alchemical free energy calculations with hybrid physics-based machine learning / molecular mechanics potentials. 2020.07.29.227959 Preprint at 10.1101/2020.07.29.227959 (2020).

40. Gapsys, V. et al. Large scale relative protein ligand binding affinities using non-equilibrium alchemy. Chem Sci 11, 1140–1152 (2019).

41. Eastman, P. et al. OpenMM 7: Rapid development of high performance algorithms for molecular dynamics. PLoS Comput Biol 13, e1005659 (2017).

42. Noske, G. D. et al. A Crystallographic Snapshot of SARS-CoV-2 Main Protease Maturation Process. J Mol Biol 433, 167118 (2021).

43. Winter, G. et al. DIALS: implementation and evaluation of a new integration package. Acta Crystallogr D Struct Biol 74, 85–97 (2018).

44. Winter, G. et al. How best to use photons. Acta Crystallogr D Struct Biol 75, 242–261 (2019).

45. Krojer, T. et al. The XChemExplorer graphical workflow tool for routine or large-scale protein-ligand structure determination. Acta Crystallogr D Struct Biol 73, 267–278 (2017).

46. Pearce, N. M. et al. A multi-crystal method for extracting obscured crystallographic states from conventionally uninterpretable electron density. Nat Commun 8, 15123 (2017).

47. Emsley, P., Lohkamp, B., Scott, W. G. & Cowtan, K. Features and development of Coot. Acta Crystallogr D Biol Crystallogr 66, 486–501 (2010).

48. Long, F. et al. AceDRG: a stereochemical description generator for ligands. Acta Crystallogr D Struct Biol 73, 112–122 (2017).

49. Laporte, M. et al. A coronavirus assembly inhibitor that targets the viral membrane protein. Nature 640, 514–523 (2025).

50. Jochmans, D., Leyssen, P. & Neyts, J. A novel method for high-throughput screening to quantify antiviral activity against viruses that induce limited CPE. Journal of Virological Methods 183, 176–179 (2012).

51. White, K. M. et al. Plitidepsin has potent preclinical efficacy against SARS-CoV-2 by targeting the host protein eEF1A. Science 371, 926–931 (2021).

52. Lombardo, F. et al. Comprehensive assessment of human pharmacokinetic prediction based on in vivo animal pharmacokinetic data, part 1: volume of distribution at steady state. J Clin Pharmacol 53, 167–177 (2013).

53. Lombardo, F. et al. Comprehensive assessment of human pharmacokinetic prediction based on in vivo animal pharmacokinetic data, part 2: clearance. J Clin Pharmacol 53, 178–191 (2013).

