## Supplementary for "Open-science discovery of DNDI-6510, a compound that addresses genotoxic and metabolic liabilities of the COVID Moonshot SARS-CoV-2 Mpro lead inhibitor"

| [1]Compound | SARS-CoV-2 Mpro $IC_{50}$ (µM) | TA-98 | | TA-100 | | TA-1535 | | TA-1537 | |
|---|---|---|---|---|---|---|---|---|---|
| | | -S9 | +S9 | -S9 | +S9 | -S9 | +S9 | -S9 | +S9 |
| **(*S*)-x38 DNDI-6510** | 0.040 | neg | neg | neg | neg | neg | neg | neg | neg |
| **(*S*)-x37 DNDI-6515** | 0.143 (rac) | neg | neg | neg | neg | neg | neg | neg | neg |
| **(*S*)-x31 DNDI-6512** | 0.074 | neg | neg | neg | neg | neg | neg | neg | neg |

**Table S1: AMES analysis of the final lead candidates (*S*)-x38, (*S*)-x37 and (*S*)-x31 against TA-98, TA-100, TA-1535 and TA-1537 strains, demonstrating the abrogation of the AMES risk due to cyclisation of the lead compounds.**

| Predicted Parameter | Method and Rationale | Value |
|---|---|---|
| **$CL_p$ (mL/min/kg)** | Mean of 4 methods | 5.17 |
| **$V_{ss}$ (L/kg)** | Geo mean of 3 most predictive allometric methods available | 1.07 |
| **Oral F (%)** | Mean of observed %F in preclinical species (mouse, rat, dog) | 62 |
| **$k_a$ ($hr^{-1}$)** | $k_a=1/MAT=1/(MRT_{PO}-MRT_{IV})$, based on mouse, dog | 0.9 |
| **$t_{1/2}$ (hrs)** | $t_{1/2} = \ln2*V_{dss}/CL$, assuming 1-cpt | 2.39 |
| **Dose (mg)*** | $Dose = (C_{min,ss}*V_{dss}*(k_a-k_e))/(F*k_a*((e^{-ke\tau})/(1- e^{-ke\tau}))-(e^{-ka\tau}/(1- e^{-ka\tau})))$ | **297 mg TID** |
| **$C_{max,ss}$ (μM)** | $C_{max,ss} = (F * e^{-ke\ Tmax} * dose)/V_{ss}$, where $T_{max} = (\ln(k_a/k_e))/(k_a-k_e)$ | 5.7 |
| **$T_{max}$ (hrs)** | $T_{max} = \ln(k_a/k_e)/(k_a-k_e)$ | 1.9 |
| **$AUC_{0-\tau,ss}$ (μM*hr)** | $AUC_{ss}=C_{av,ss}*\tau$ | 18 |
| **$R_{AC}$** | $R_{AC}=1/(1-e^{(-ke*t)})$ | 1.1 |
| **Dose (mg)*** | $Dose = (C_{min,ss}*V_{dss}*(k_a-k_e))/(F*k_a*((e^{-ke\tau})/(1- e^{-ke\tau}))-(e^{-ka\tau}/(1- e^{-ka\tau})))$ | **1010 mg BID** |
| **$C_{max,ss}$ (μM)** | $C_{max,ss} = (F * e^{-ke\ Tmax} * dose)/V_{ss}$, where $T_{max} = (\ln(k_a/k_e))/(k_a-k_e)$ | 10 |
| **$T_{max}$ (hrs)** | $T_{max} = \ln(k_a/k_e)/(k_a-k_e)$ | 1.9 |
| **$AUC_{0-\tau,ss}$ (μM*hr)** | $AUC_{ss}=C_{av,ss}*\tau$ | 61 |
| **$R_{AC}$** | $R_{AC}=1/(1-e^{(-ke*t)})$ | 1.0 |

**Table S2: Human PK prediction for DNDI-6510, aiming for continuous exposure above IC90 380 nM free, Human fu 0.46, target Cmin of 0.826 uM total.**

| Measured Parameter | (*S*)-x31 DNDI-6512 | (*S*)-x37 DNDI-6515 | (*S*)-x38 DNDI-6510 |
|---|---|---|---|
| **Molecular weight** | 434.927 | 483.355 | 474.948 |
| **LogD (measured) pH 7.4** | 2.4 | 2.3 | 1.7 |
| **tPSA /Å2calc** | | | |
| **pKa** | 5.5 | 3.4 | 4.5 |
| **Kinetic solubility (µM) pH 1** | 3500 | 330 | 3100 |
| **Kinetic solubility (µM) pH 6.8** | 210 | 30 | 540 |

**Table S3**: **Basic compound characteristics for (S)-x38 DNDI-6510, (S)-x37 and (S)-x31.**

| | Fraction unbound (%) | Fraction unbound (%) | Fraction unbound (%) |
|---|---|---|---|
| Species | ***S*)-x38 DNDI-6510** | **(*S*)-x31 DNDI-6512** | **(*S*)-x37 DNDI-6515** |
| **Mouse** | 28 | 16 | 15 |
| **Rat** | 54 | 53 | 22 |
| **Dog** | 12 | | 16 |
| **Cyno** | 60 | 57 | 29 |
| **Minipig** | 56 | 33 | 32 |
| **Human** | 46 | 37 | 9 |
| **FCS (2%)** | 75 | | 96 |

**Table S4: Plasma protein binding of DNDI-6510 across species, and cell culture media.**

| | ***S*)-x38 DNDI-6510** | **(*S*)-x31 DNDI-6512** | **(*S*)-x37 DNDI-6515** |
|---|---|---|---|
| **Rat** | 1.2 | 0.91 | 1.0 |
| **Human** | 0.73 | 0.67 | 0.65 |

**Blood-to-plasma ratio.**

| Readout | **(*S*)-x38 DNDI-6510** | **(*S*)-x31 DNDI-6512** | **(*S*)-x37 DNDI-6515** |
|---|---|---|---|
| **A>B (cm/sec)** | 0.5 | 2.5 | 0.9 |
| **B>A (cm/sec)** | 14.5 | 33.8 | 25 |
| **ER** | 31 | 13 | 27 |
| **A>B (Low efflux)** | 6.8 | 20.2 | 15.7 |

**MDCK permeability.**

| Compound ID (IC50) uM | **(*S*)-x31 DNDI-6512** | **(S)-x38 DNDI-6510** | **Ac-DEVD-CHO** |
|---|---|---|---|
| **Caspase-1** | >100 | >100 | 0.167 |
| **Caspase-2** | >100 | >100 | 2.32 |
| **Caspase-3** | >100 | >100 | 0.000691 |
| **Caspase-5** | >100 | >100 | 2.43 |
| **Caspase-6** | >100 | >100 | 0.0575 |
| **Caspase-7** | >100 | >100 | 0.00467 |
| **Caspase-8** | >100 | >100 | 0.00232 |
| **Caspase-9** | >100 | >100 | 0.104 |
| | | | E-64 |
| **Calpain1** | >100 | >100 | 0.067 |
| **Cathepsin B** | >100 | >100 | 0.0265 |
| **Cathepsin K** | >100 | >100 | 0.0109 |
| **Cathepsin L** | >100 | >100 | 0.0238 |
| **Cathepsin S** | >100 | >100 | 0.00501 |

**Table S5. Selectivity of DNDI-6510 against a panel of mammalian proteases.** The inhibitory activity of (S)-x31, (S)-x37 and (S)-x38 was evaluated using FRET-based assay format at several mammalian proteases up to 100 uM concentration.

| | (S)-x38 DNDI-6510 | (*S*)-x37 DNDI-6515 | (*S*)-x31 DNDI-6512 |
|---|---|---|---|
| **PSP (Eurofins): hr Al2a (Binding) (AC50)** | >30 | >30 | >30 |
| **PSP (Eurofins): r Bzd (Binding) (AC50)** | >30 | >30 | >30 |
| **PSP (Eurofins): hr D1 (Binding) (AC50)** | >30 | >30 | >30 |
| **PSP (Eurofins): hr D3 (Binding) (AC50)** | >30 | >30 | >30 |
| **PSP (Eurofins): hr DAT (Binding) (AC50)** | >30 | >30 | >30 |
| **PSP (Eurofins): hr M1 (Binding) (AC50)** | >30 | >30 | >30 |
| **PSP (Eurofins): hr M2 (Binding) (AC50)** | >30 | >30 | >30 |
| **PSP (Eurofins): hr DOP (Binding) (AC50)** | >30 | >30 | >30 |
| **PSP (Eurofins): hr MOP (Binding) (AC50)** | >30 | >30 | >30 |
| **PSP (Eurofins): hr 5HT1A (Binding) (AC50)** | >30 | >30 | >30 |
| **PSP (Eurofins): hr MAO (Antagonism) (AC50)** | >30 | >30 | >30 |
| **PSP (Eurofins): hr NET (Binding) (AC50)** | >30 | >30 | >30 |
| **PSP (Eurofins): hr ACES (Antagonism) (AC50)** | >30 | >30 | >30 |
| **PSP (Eurofins): hr 5HTT (Binding) (AC50)** | >30 | >30 | >30 |
| **PSP (Eurofins) - hr Ad1 (Binding) (AC50)** | >30 | >30 | >30 |
| **PSP (Eurofins): hr AR (Binding) (AC50)** | >30 | 10 | >30 |
| **PSP (Eurofins): hr H3 (Binding) (AC50)** | >30 | >30 | >30 |
| **PSP (Eurofins): hr TP (Agonism) (AC50)** | >30 | >30 | >30 |
| **PSP (Eurofins): hr TP (Antagonism) (AC50)** | >30 | 10 | >30 |
| **PSP (Eurofins): hr Al1a - (Binding) (AC50)** | >30 | >30 | >30 |
| **PSP (Eurofins): hr PR (Binding) (AC50)** | >30 | >30 | >30 |
| **PSP (Eurofins): hr 5HT2B (Agonism) (AC50)** | >30 | >30 | >30 |
| **PSP (Eurofins): h Thrombin (Antagonism) (AC50)** | >30 | >30 | >30 |
| **PSP (Eurofins): hr PDE3A (Antagonism) (AC50)** | >30 | >30 | >30 |
| **PSP (Eurofins): hr PDE4D2 (Antagonism) (AC50)** | >30 | >30 | >30 |
| **PSP (Eurofins): hr ERalpha (Binding) (AC50)** | >30 | >30 | >30 |
| **PSP (Eurofins): hr COX1 (Antagonism) (AC50)** | >30 | >30 | >30 |
| **PSP (Eurofins): hr COX2 (Antagonism) (AC50)** | >30 | >30 | >30 |
| **PSP (Eurofins): HEK Native Antagonism (AC50)** | >30 | 14.01 | >30 |

**Table S6: Eurofins off target panel**

| | (*S*)-x38 DNDI-6510 | (*S*)-x31 DNDI-6512 | (*S*)-x37 DNDI-6515 |
|---|---|---|---|
| **hERG Binding $IC_{50}$ µM** | >30 | >30 | *>30* |
| **hERG PC $IC_{50}$ µM** | >30 | >30 | 20 |
| **Nav1.5 $IC_{50}$ µM** | >50 | >50 | >50 |
| **BSEP $IC_{50}$ µM** | 45 | 41 | 54 |
| **Principal panel (31 targets) $IC_{50}$ µM** | All >30 | All >30 | All >30 |
| **Kinase safety panel (59 kinases) $IC_{50}$ µM** | All >30 | All >30 | All >30 |
| **Comprehensive panel (77 panel) % inhib @ 10uM** | Clean | Clean | Mt2 (63%)<br>AdT (57%) |

**Table S7: Ion channels.**

| Compound | Hepatic Microsomes | t1/2 (min) | CLint (µl/min/mg) |
|---|---|---|---|
| *S*)-x38 DNDI-6510 | Mouse | 74.4 | 23.2 |
| | Rat | 108.4 | 16 |
| | Dog | 92.2 | 18.8 |
| | Minipig | 122.2 | 14.2 |
| | Cyno | 19.3 | 90 |
| | Human | 244.4 | 7.1 |
| | 8HUM | 100.1 | 14 |
| (*S*)-x31 DNDI-6512 | Mouse | 12.1 | 143.2 |
| | Rat | 16.4 | 105.5 |
| | Dog | 38.7 | 44.9 |
| | Minipig | 7.9 | 219.5 |
| | Cyno | 4.2 | 412.3 |
| | Human | 27.9 | 62.2 |
| | 8HUM | 68.6 | 20 |
| (*S*)-x37 DNDI-6515 | Mouse | 12.6 | 137.8 |
| | Rat | 23.4 | 74.2 |
| | Dog | 36.7 | 47.3 |
| | Minipig | 11.6 | 149.6 |
| | Cyno | 4.4 | 389.8 |
| | Human | 37.6 | 46.1 |
| | 8HUM | 68.4 | 20 |

**Table S8. In vitro metabolic stability of DNDI-6510 in hepatic microsomes from preclinical species and human.** Metabolic stability was examined in hepatic microsomes using compounds at a concentration of 1 µM. Incubations were conducted in duplicate and mean stability data (half-life (t1/2) and intrinsic clearance (CLint) is depicted. Incubations with 8HUM microsomes were carried out in singlicate with compounds at a final concentration of 0.5 uM.

| ***Hepatocytes (Novartis)*** | (*S*)-x38 DNDI-6510 | (*S*)-x31 DNDI-6512 | (*S*)-x37 DNDI-6515 |
|---|---|---|---|
| **Cyno** | 16 | 46 | *41* |
| **Dog** | 4 | 9 | *6* |
| **Rat** | 4 | 32 | *43* |
| **Minipig** | 4 | 47 | *44* |
| **Human** | 4 | 20 | *7* |

| ***Hepatocyte (Cyprotex)*** | **(S)-x38** DNDI-6510 | **(S)-x31** DNDI-6512 | **(*S*)-x37** DNDI-6515 |
|---|---|---|---|
| **Mouse** | 15.4 | 87.1 | 117.2 |
| **Rat** | <4 | 35.8 | 24.7 |
| **Dog** | 4.7 | 7.1 | 5.9 |
| **Monkey** | 7.9 | 46.1 | 40.6 |
| **Minipig** | <4 | 52.9 | 44.2 |
| **Human** | <4 | 6.6 | 7.3 |

**Table S9. In vitro metabolic stability of DNDI-6510 in hepatocytes from preclinical species and human.** Metabolic stability was examined in intestinal microsomes using test compounds at a concentration of 1 μM. Incubations were conducted in duplicate and mean stability data (half-life (t1/2) and intrinsic clearance (CLint) is depicted.

| **Viral strain** | **Cell type** | ***S*)-x38** DNDI-6510 | **(*S*)-x37** DNDI-6515 | **(*S*)-x31** DNDI-6512 | **Nirmatrelvir** | **Ensitrelvir** |
|---|---|---|---|---|---|---|
| **229E** | Huh7 | **>50** | **>50** | **>50** | | |
| **GHB** | VeroE6 | >10 | >10 | >10 | >10 | >10 |
| **B1.1.7** | A549 | >10 | >10 | >10 | >10 | >10 |

**Table S10. Cytotoxicity shown as $CC_{50}$ values, observed for compounds tested in recombinant cell lines used for antiviral assays.**

| | | IC50 of compounds in Mpro and CatB enzymatic assays (uM) | | | | | | | | EC50 (CPE) | |
|---|---|---|---|---|---|---|---|---|---|---|---|
| **#** | **CPD ID** | **SARS-2** | **229E** | **OC43** | **MERS** | **SARS** | **HKU1** | **NL63** | **CatB** | **229E** | **OC43** |
| **1** | **(*S*)-x31 DNDI-6512** | 0.165 | > 20 | 7.783 | > 20 | 0.337 | 5.736 | > 20 | > 20 | >20 | 10.40 |
| **2** | **(*S*)-x37 DNDI-6515** | 0.361 | > 20 | > 20 | > 20 | 0.550 | > 20 | > 20 | > 20 | >20 | >20 |
| **3** | **(*S*)-x38 DNDI-6510** (rac) | 0.171 | > 20 | > 20 | > 20 | 0.323 | > 20 | > 20 | > 20 | >20 | >20 |
| **Ref.** | GC376 | 0.026 | 0.123 | 0.042 | 0.267 | 0.042 | 0.017 | 0.139 | > 50 | NA | NA |
| **Ref.** | Leupeptin hemisulfate salt | NA | NA | NA | NA | NA | NA | NA | 0.057 | NA | NA |
| **Ref.** | Remdesivir | NA | NA | NA | NA | NA | NA | NA | NA | 0.086 | 0.018 |

**Table S11. Inhibition of human coronavirus main proteases.** Test compounds were assessed for its ability to inhibit the proteolytic activity of MPro of SARS-CoV-2 and related coronaviruses in a FRET assay up to 20 μM. Data is given as IC50 measurements (uM).

| | **DNDI-6510 50 mg/kg** | **DNDI-6510 15 mg/kg** | **DNDI-6510 50 mg/kg** | **DNDI-6510 150 mg/kg** | **DNDI-6510 50 mg/kg** | **DNDI-6510 150 mg/kg** |
|---|---|---|---|---|---|---|
| | WT | 50 mg/kg ABT | | | 8HUM | |
| $C_{max}$ (ng/mL) | 5,987 | 4,617 | 7,210 | 26,000 | 4,124 | 24,004 |
| $T_{max}$ (h) | 0.5 | 1 | 2 | 0.5 | 1 | 1 |
| $AUC_{last}$ (ng.h/mL) | 9,623 | 17,777 | 41,887 | 146,021 | 1,002,298 | 7,125,320 |
| $AUCM_{last}$ (ng.h.h/mL) | 13,501 | | 169,220 | | | |
| Cl_F_obs (mL/min/kg) | 90 | 13 | 20 | 17 | 48 | 21 |
| T ½ elim (h) | 2.5 | 2.5 | 1.5 | 1.4 | | |
| $MRT_{last}$ (h) | 1.4 | 3.7 | 4 | 4.2 | 2.9 | 249.7 |
| F | 39% | | | | | |

**Table S12. Pharmacokinetic assessment and analysis of (S)-x38 DNDI-6510 in wild-type mice, in wild-type mice with ABT co-dosing, and in a humanized mouse model (8HUM).**

| Species | Formulation | Dose (mg/kg) | $C_{max}$ (ng/ml) | $T_{max}$ (h) | $AUC_{last}$ (ng·h/ml) | CLp (ml/min/kg) | Vdss (l/kg) | t1/2 (h) | Oral F (%) |
|---|---|---|---|---|---|---|---|---|---|
| **Rat #** | [V] | 1 (IV) | | | 661 ± 42.5 | 25 | 1.4 | 0.86 | |
| **Rat #** | [X] | 2 (PO) | 147 | 1 | 428 ± 20.6 | | | | 32 |
| **Rat #** | [X] | 5 (PO) | 76 | 1 | 594 | | | | 18 |
| **Rat #** | [X] | 10 (PO) | 195 | 1 | 762 | | | | 11 |
| **Rat ~** | [Y*] | 10 (PO) | 579 | 1 | 2469 | 72 | | 2 | 37 |
| **Rat ~** | [Z] | 30 (PO) | 1052 | 2 | 3946 | 130 | | 2.7 | 20 |
| **Rat ~** | [Z] | 60 (PO) | 2270 | 2 | 14284 | 70 | | 2.8 | 36 |
| **Dog #** | [W] | 1 (IV) | | | 586 | 23.5 | 1.4 | 0.86 | |
| **Dog #** | [X] | 2 (PO) | 924 | 1 | 2480 | | | | >100 |
| **Dog #** | [X] | 5 (PO) | 2100 | 1 | 4150 | | | | >100 |
| **Dog #** | [X] | 10 (PO) | 3550 | 1 | 7940 | | | | >100 |

**Table S13. Preclinical pharmacokinetics of SARS-CoV2 Mpro inhibitor (S)-x38 DNDI-6510.** All rat pharmacokinetics were conducted in male gender, and in the fed state. Pharmacokinetic parameters were calculated from plasma concentration–time data and are reported as mean (± S.D. for n=3). For IV studies at 1 mg/kg, (S)-x38 was administered as 20% w/w propylene glycol (PG), 10% w/w Cremophor RH40 and 70% w/w sterile water in rat [V], and as 5% w/w DMSO and 95% w/w saline [W] in dog. For po studies in rats, (S)-x38 DNDI-6510 was administered as a suspension, for 2, 5 and 10 mg/kg in 2.5%w/v Vit E TPGS in sterile water [X], and for 30 and 60 mg/kg in 1 % HPMC in water with 1 % tween 80 (suspension) [Z], or as a solution for the racemate dosing at 10 mg/kg (*) in 10% DMSO, 5% Tween 80, 30% PEG400, 10% PG, 45% Saline (solution) [Y].
### Aptuit, ~TCG, * on racemate

| CYP enzyme | Inhibition | TDI (-NADPH) | | TDI (+NADPH) | | Fold shift | Reaction phenotyping | |
|---|---|---|---|---|---|---|---|---|
| | IC50 (µM) | IC50 (µM) | SE (µM) | IC50 (µM) | SE (µM) | | t1/2 (min) | SE t1/2 (min) |
| **CYP3A4** | | | | | | | 155 | 52.3 |
| **CYP3A4 (testosterone)** | >25 | >25 | | >25 | | | | |
| **CYP3A4 (midazolam)** | >25 | >25 | | 20.6 | 0.752 | >1.21 | | |
| **CYP2C19** | >25 | >25 | | >25 | | | 547 | 498 |
| **CYP1A2** | >25 | >25 | | >25 | | | -1250 | 2280 |
| **CYP2C8** | >25 | >25 | | >25 | | | 704 | 1140 |
| **CYP2D6** | >25 | >25 | | >25 | | | -5540 | 23900 |
| **CYP2B6** | >25 | >25 | | >25 | | | 368 | 412 |
| **CYP2C9** | >25 | >25 | | >25 | | | -2020 | 4490 |

**Table S14: Cytochrome P450 (CYP) enzymes activity of** (S)**-x38 DNDI-6510.** (S)-x38 DNDI-6510 was tested (n=7) for enzyme inhibition, and time-dependent inhibition (TDI) with and without NADPH after 30 minutes of incubation. For the reaction phenotyping analysis, the amount of DNDI-6510 was determined after incubation with seven clinically relevant CYP enzyme, and amount of non-metabolised compound determined at 5, 15, 30, and 45 minutes.

| | | | *S*)-x38 DNDI-6510 | |
|---|---|---|---|---|
| **Species** | Nuclear Hormone receptor (NHR) | EC50/uM | Max Activation-fold | Max Activation conc/uM |
| **Human** | AhR | | 1.19 | 100 |
| | CAR1 | | 1.83 | 100 |
| | CAR3 | 15.5 | 1.03 | 0.300 |
| | PXR | | 16.6 | 100 |
| **Dog** | AhR | | 1.22 | 100 |
| | CAR3 | | 1.09 | 30.0 |
| | PXR | | 9.96 | 100 |
| **Rat** | AhR | | 1.20 | 100 |
| | CAR3 | | 1.15 | 100 |
| | PXR | | 10.5 | 100 |
| **Mouse** | AhR | | 2.22 | 100 |
| | CAR1 | | 1.44 | 30.0 |
| | PXR | 4.71 | 4.66 | 30.0 |

**Table S15: Nuclear Hormone Receptor (NHR) induction profile of (S)-x38 DNDI-6510 across species.**

| **Solubility mg/ml @ 2 hours** | **Water** | **pH 7.4 buffer** | **FaSSIF pH 6.5** | **FeSSIF pH 5** | **FaSGIF pH 1.6** |
|---|---|---|---|---|---|
| **Form 1** | **0.02** | **0.01** | **0.03** | **0.09** | **0.31** |
| HCL salt | **0.46** | **0.02** | **0.03** | **0.04** | **0.89** |
| Amorphous | **0.19** | **0.62** | **0.23** | **0.38** | **1.23** |
| Form 11 | **0.06** | **0.04** | **0.23** | **0.34** | **2.34** |
| Hydrate | **0.01** | **0.01** | **0.01** | **0.02** | **0.49** |

**Table S15**: (S)-x38 DNDI-6510 Solid state analysis. All forms – after 2 hrs, [ ] falls as Form 2 hydrate comes out of solution

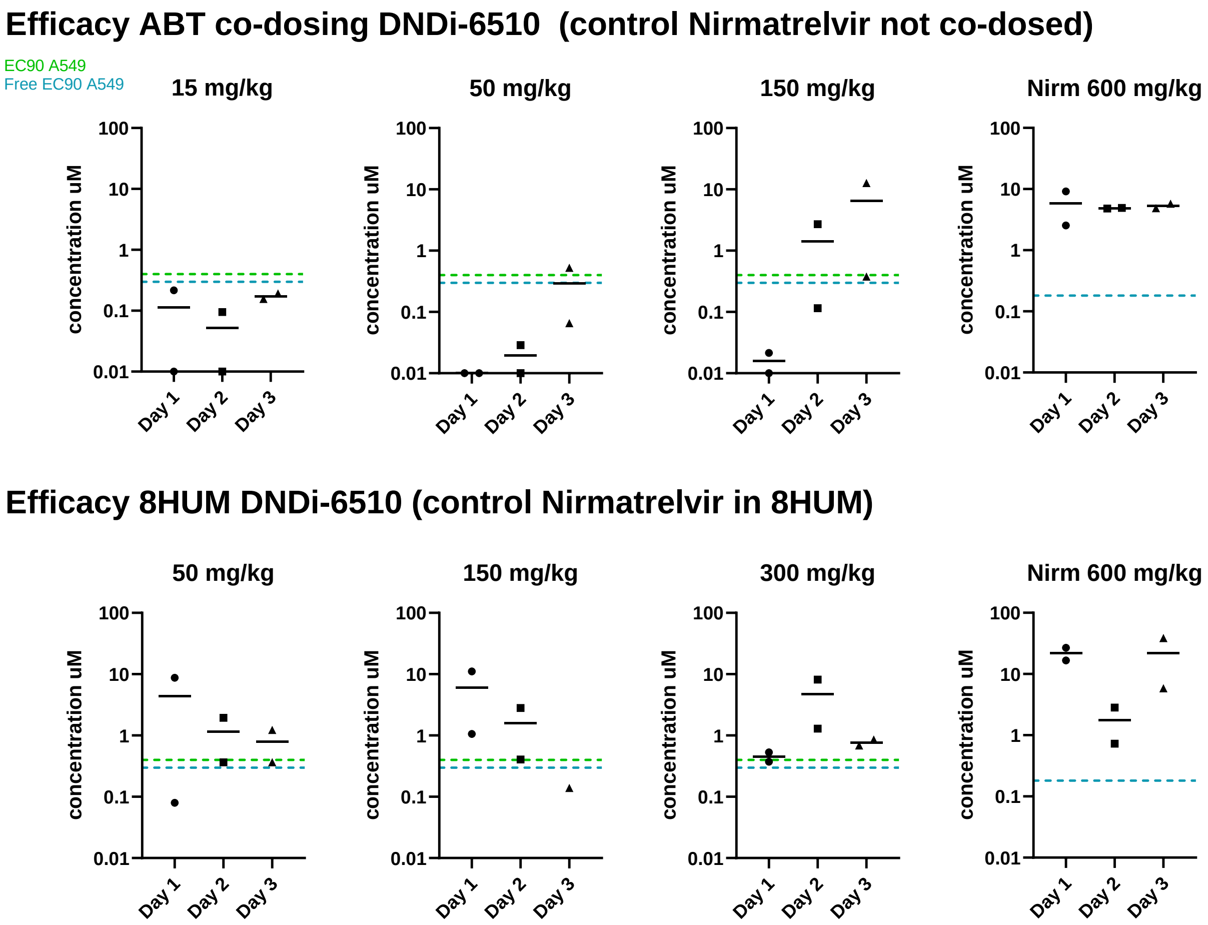

**Figure S1: Pharmacokinetic exposure of (S)-x38 DNDI-6510 in satellite animals dosed alongside the mouse efficacy experiment.**

**A**

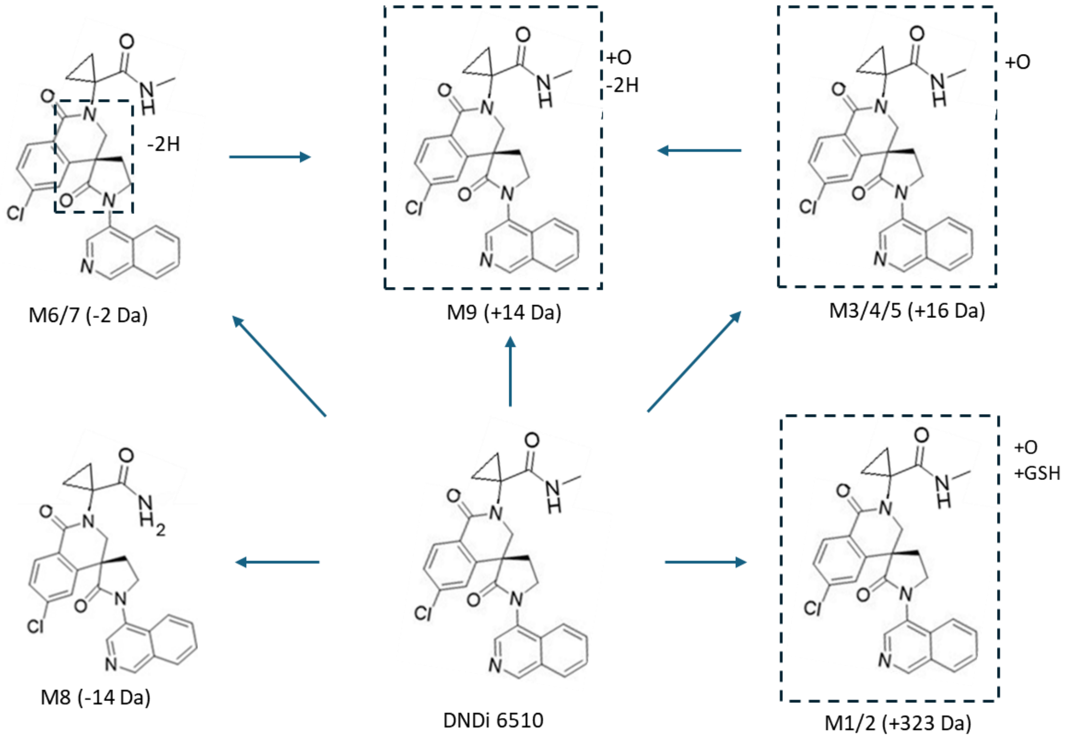

**B**

| Species | Human | Rat | Mouse | Dog | Monkey | Minipig |
|---|---|---|---|---|---|---|
| DNDI-6510 | 0.997 | 0.914 | 0.536 | 0.691 | 0.911 | 0.953 |
| M1 +323 | | 0.020 | 0.157 | 0.170 | | 0.007 |
| M2 +323 | | | | 0.062 | | |
| M3 +16 | | 0.010 | 0.009 | 0.033 | | 0.002 |
| M4 +16 | | 0.012 | 0.006 | 0.037 | | 0.005 |
| M5 +16 | | 0.008 | 0.017 | | 0.016 | 0.009 |
| M6 -2 | | 0.010 | | 0.004 | | 0.005 |
| M7 -2 | 0.001 | 0.017 | 0.040 | 0.003 | 0.033 | 0.018 |
| M8 -14 | | 0.001 | 0.103 | | 0.013 | 0.001 |
| M9 +14 | 0.002 | 0.009 | 0.131 | 0.027 | | |

**Figure S2: In vitro metabolite identification of (S)-x38 DNDI-6510.** To identify in vitro metabolites, compounds were incubated with 1 million cryopreserved hepatocytes from different species at a concentration of 1 $\mu$M in 96 well plates, with 1000 rpm for 1, 10, 20, 40, 60 and 80 min*,* respectively. Clearance of DNDI-6510 in cryopreserved hepatocytes is detailed in Table S9. (A) Metabolites identified in hepatocyte assays, identified in LC-MS through a full scan with data-dependent MS/MS. LC separation using a generic gradient up to 10 min using a Q-Exactive, and data analysed through SSID. (B) Cross-species overview and quantification of identified metabolites.

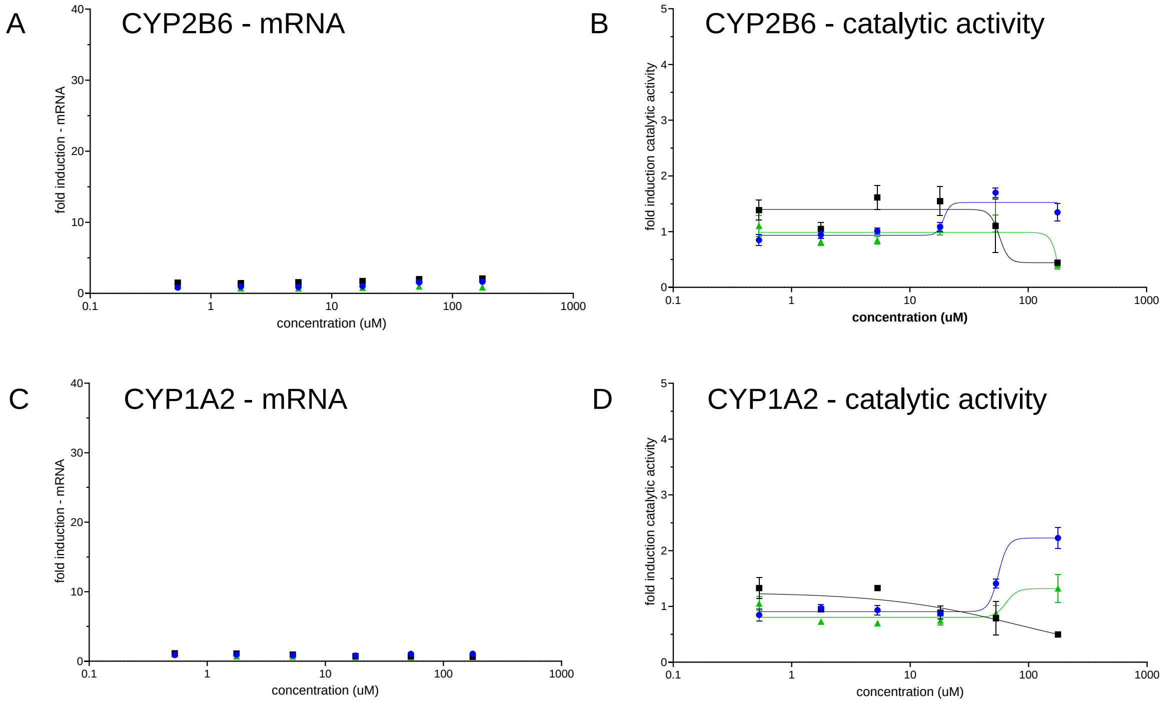

**Figure S3: CYP enzyme induction in cryopreserved hepatocytes in response to (S)-x38 DNDI-6510.** No metabolic induction were measured for (A) CYP2B6 mRNA, (B) CYP2B6 catalytic activity or (C) CYP1A2 mRNA or (D) CYP1A2 catalytic activity after incubation with increasing concentrations of (S)-x38 DNDI-6510 up to 177 uM.
